# Synthesis and Characterization of ULK1/2 Kinase Inhibitors that Inhibit Autophagy and Upregulate Expression of Major Histocompatibility Complex I for the Treatment of Non-Small Cell Lung Cancer

**DOI:** 10.1101/2025.09.05.674519

**Authors:** Fabiana Izidro A. L. Layng, Huiyu Ren, Nicole A. Bakas, Dhanya R. Panickar, Lester J. Lambert, Maria Celeridad, Jiaqian Wu, Laurent De Backer, Preeti Chandrachud, Allison S. Limpert, Mitchell Vamos, Apirat Chaikuad, Betsaida B. Verdugo, Patrick M. Hagan, Sonja N. Brun, Lutz Tautz, Stefan Knapp, Reuben J. Shaw, Guy S. Salvesen, Douglas J. Sheffler, Nicholas D. P. Cosford

## Abstract

Autophagy inhibition represents a promising therapeutic approach for the management of various cancers including non-small cell lung cancer (NSCLC). We previously reported **SBP-7455**, a dual inhibitor of unc-51-like kinase 1 (ULK1) and its homologue ULK2, and described its effects on triple-negative breast cancer (TNBC) cells. Herein we report the design, synthesis, and characterization of **SBP-5147** and **SBP-7501**, two new dual ULK1/2 inhibitors that are cytotoxic against NSCLC cells, inhibit autophagic flux in A549 cells, and present greater oral exposure than **SBP-7455** at a lower dose. In addition, **SBP-5147** effectively modulates autophagy and increases the expression of major histocompatibility complex (MHC) class I in NSCLC cells, which may support the rationale for ULK1/2 inhibition as a strategy to overcome resistance to immunotherapy. Together these data support the use of ULK inhibitors as part of a cancer treatment strategy, both as a single agent as well as in combination with current therapies.

## INTRODUCTION

Macroautophagy (hereafter referred to as autophagy) is a lysosome-dependent, highly conserved physiological process in eukaryotic cells characterized by intracellular degradation and recycling, which is induced by stress stimuli to maintain cellular homeostasis and support cell survival ^1, 2^. The autophagic system consists of a multistep pathway mediated by 37-autophagy-related proteins (ATGs) that are responsible for transporting cytoplasmic cargo to the lysosome for degradation ^3^. Autophagy is regulated by the ULK1 preinitiation complex which includes the unc-51-like kinase 1 (ULK1) or its functional homologue ULK2 (ULK1/2), ATG13, RIB-inducible coiled-coil protein 1 (FIP200), and ATG101. Under physiological conditions, the mechanistic target of rapamycin (mTOR) inhibits autophagy by phosphorylating ULK1 at S757, S638, and S758 residues. However, during cellular stress, AMP-activated protein kinase (AMPK) inhibits mTOR, allowing for the dephosphorylation of ULK1 at inhibitory serine residues, and promoting the phosphorylation of ULK1 at S555, S777, S317, and S467, thus triggering the activity of the ULK1 complex and inducing autophagy ^2–4^. Activation of the ULK1 complex stimulates phagophore nucleation via phosphorylation of Beclin-1 and vacuolar protein sorting 34 (Vps34), a class III phosphatidylinositol 3-kinase (PI3K) ^3, 5^. Additionally, ULK1 regulates the trafficking of ATG9, a transmembrane protein essential for autophagosome maturation, by phosphorylating the protein at S14, thus allowing for its relocation from the plasma membrane to the developing autophagosome ^6^. As ULK1 plays a central role in autophagy initiation and is the only serine/threonine (S/T) kinase in the core autophagy pathway, it is a promising target for drug development. Compounds that selectively inhibit autophagy have demonstrated clinical relevance in the context of cancer treatment ^2, 7^. In addition to our published pyrimidine-based compounds, **SBI-0206965** and **SBP-7455** (**Figure S1**), several other ULK1/2 inhibitors targeting autophagy in cancer have been described, including MRT68921 and MRT67307, 2-aminopyrimidine derivatives originally synthesized as TBK1 inhibitors ^8^ that block autophagosome maturation ^7–9^; the pyrazole-derived ULK-101, which suppresses the nucleation of autophagic vesicles and autophagic turnover ^10^; hesperidin, a bioflavonoid that induces apoptosis and cell cycle arrest in cancer cells and potently binds to ULK1/2 ^9, 11–13^; and DCC-3116, an orally available dual ULK1/2 inhibitor that prevents autophagosome formation and lysosomal degradation ^8^. The latter is currently in early-phase clinical trials in patients with advanced solid tumors presenting RAS/MAPK pathway mutations (NCT04892017).

The role of autophagy in cancer can vary due to the biological characteristics of the tumor, the stage of the cancer, and what treatment strategies have been employed. In some early cancers, autophagy may function as a tumor suppressor by preserving genomic integrity and preventing proliferation and inflammation ^14^. In more advanced cancers, autophagy promotes tumor progression by enabling cancer cell proliferation and survival in metabolically stressed and treatment-challenged tumor microenvironments, contributing to cancer progression and resistance to treatment ^14, 15^.

In non-small cell lung cancer (NSCLC), which accounts for 85% of all lung cancer cases, the identification of targetable genetic alterations and use of immune checkpoint inhibitors (ICI) have propelled precision oncology strategies, resulting in extraordinary survival benefits in selected patients ^16, 17^. However, the impact of personalized medicine on overall survival rates for NSCLC has been modest, particularly in advanced and metastatic disease ^17^, underlining the need for development of therapeutic approaches that tackle biological processes that promote tumor progression and influence treatment responses, such as autophagy ^1, 18^. To this end, preclinical studies have linked the upregulation of autophagy with resistance to Osimertinib, a standard-of-care, third-generation epidermal growth factor receptor (EGFR)-tyrosine kinase inhibitor (TKI) for *EGFR*-mutant NSCLC ^19^. ULK1 has been shown to be overexpressed in lung cancer cell lines such as A549 and HCC827, and its upregulation correlates with poor prognosis in NSCLC patients ^20, 21^. Moreover, ULK1 inhibition by **SBI-0206965**, our first-generation ULK1 chemical probe, synergizes with cisplatin against NSCLC cells and re-sensitizes cisplatin-resistant cells ^21^. ULK1 inhibition has also shown synergy with immune checkpoint blockade, enhancing antitumor immunity, and resulting in tumor regression in murine models of *LKB1*-mutant NSCLC ^22^.

Thus, to support strategies that could improve treatment responses in NSCLC, herein we report **SBP-5147** and **SBP-7501** (**Table 1**), novel dual ULK1/2 small-molecule inhibitors developed from a structure-based rational design that are more potent than our previously published chemical probe, **SBP-7455** ^2^. Both **SBP-5147** and **SBP-7501** exhibit improved biochemical potency, increased intracellular binding to ULK1/2, and strong cytotoxicity as single agents against NSCLC cells. In addition, **SBP-5147** successfully inhibited autophagy in vitro and increased the expression of major histocompatibility complex (MHC) class I in NSCLC cells.

**Table 1.**
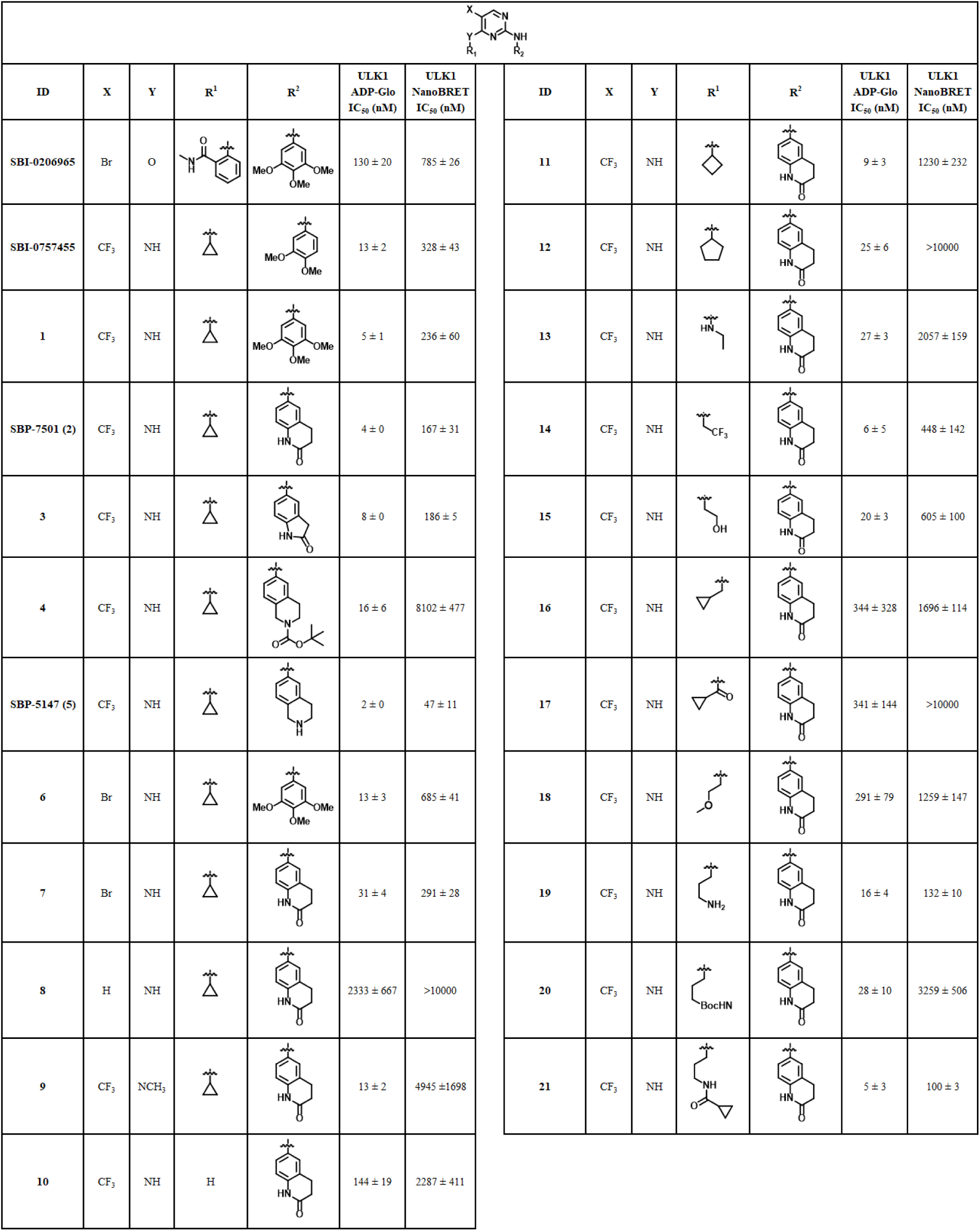
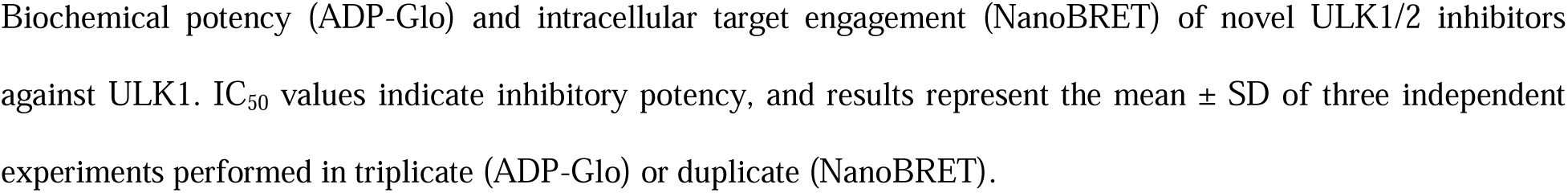
Structures of ULK1/2 inhibitors and ULK1 ADP-Glo and NanoBRET IC_50_ values.

|  |  |  |  |  |  |  |  |  |  |  |  |  |  |
| --- | --- | --- | --- | --- | --- | --- | --- | --- | --- | --- | --- | --- | --- |
| ID | X | Y | R<sup>1</sup> | R<sup>2</sup> | ULK1<br>ADP-Glo<br>IC<sub>50</sub> (nM) | ULK1<br>NanoBRET<br>IC<sub>50</sub> (nM) | ID | X | Y | R<sup>1</sup> | R<sup>2</sup> | ULK1<br>ADP-Glo<br>IC<sub>50</sub> (nM) | ULK1<br>NanoBRET<br>IC<sub>50</sub> (nM) |
| SBI-0206965 | Br | O |  |  | 130 ± 20 | 785 ± 26 | 11 | CF<sub>3</sub> | NH |  |  | 9 ± 3 | 1230 ± 232 |
| SBI-0757455 | CF<sub>3</sub> | NH |  |  | 13 ± 2 | 328 ± 43 | 12 | CF<sub>3</sub> | NH |  |  | 25 ± 6 | >10000 |
| 1 | CF<sub>3</sub> | NH |  |  | 5 ± 1 | 236 ± 60 | 13 | CF<sub>3</sub> | NH |  |  | 27 ± 3 | 2057 ± 159 |
| SBP-7501 (2) | CF<sub>3</sub> | NH |  |  | 4 ± 0 | 167 ± 31 | 14 | CF<sub>3</sub> | NH |  |  | 6 ± 5 | 448 ± 142 |
| 3 | CF<sub>3</sub> | NH |  |  | 8 ± 0 | 186 ± 5 | 15 | CF<sub>3</sub> | NH |  |  | 20 ± 3 | 605 ± 100 |
| 4 | CF<sub>3</sub> | NH |  |  | 16 ± 6 | 8102 ± 477 | 16 | CF<sub>3</sub> | NH |  |  | 344 ± 328 | 1696 ± 114 |
| SBP-5147 (5) | CF<sub>3</sub> | NH |  |  | 2 ± 0 | 47 ± 11 | 17 | CF<sub>3</sub> | NH |  |  | 341 ± 144 | >10000 |
| 6 | Br | NH |  |  | 13 ± 3 | 685 ± 41 | 18 | CF<sub>3</sub> | NH |  |  | 291 ± 79 | 1259 ± 147 |
| 7 | Br | NH |  |  | 31 ± 4 | 291 ± 28 | 19 | CF<sub>3</sub> | NH |  |  | 16 ± 4 | 132 ± 10 |
| 8 | H | NH |  |  | 2333 ± 667 | >10000 | 20 | CF<sub>3</sub> | NH |  |  | 28 ± 10 | 3259 ± 506 |
| 9 | CF<sub>3</sub> | NCH<sub>3</sub> |  |  | 13 ± 2 | 4945 ± 1698 | 21 | CF<sub>3</sub> | NH |  |  | 5 ± 3 | 100 ± 3 |
| 10 | CF<sub>3</sub> | NH | H |  | 144 ± 19 | 2287 ± 411 |  |  |  |  |  |  |  |

Biochemical potency (ADP-Glo) and intracellular target engagement (NanoBRET) of novel ULK1/2 inhibitors against ULK1. IC<sub>50</sub> values indicate inhibitory potency, and results represent the mean ± SD of three independent experiments performed in triplicate (ADP-Glo) or duplicate (NanoBRET).

## RESULTS AND DISCUSSION

To optimize and further characterize more potent and selective small molecule ULK1/2 inhibitors that are cytotoxic against cancer cells, we applied our previously reported testing platform ^2^ that prioritizes metrics for both biochemical and cellular efficacy. To measure the efficacy of the novel chemical probes as biochemical inhibitors of ULK1/2, we used an in vitro endpoint, high-throughput formatted ADP-Glo^TM^ kinase assay (Promega) ^11, 23^. This assay is carried out in two steps after the kinase reaction, including depletion of remaining ATP and conversion of enzyme reaction product ADP to ATP combined with a luciferase/luciferin reaction. Therefore, the kinase activity is measured by luminescence, which is proportional to the amount of ADP generated during the kinase reaction ^23^ (**Figure S2**, **A**).

To quantify intracellular target engagement, we employed a bioluminescence energy transfer NanoBRET assay, optimized for ULK1/2 and compatible with a high-throughput workflow. This approach utilizes live cells expressing ULK1 or ULK2 tagged with a 19 kDa luciferase domain (NanoLuc) and a cell-permeable bivalent fluorescent energy transfer (FRET) probe that emits a fluorescent signal ^2, 24, 25^. Bioluminescence resonance energy transfer (BRET) is attained inside intact cells by reversible binding of the fluorescent tracer to the intracellular target ^25^. ULK1/2 inhibitors that penetrate the cells compete with the tracer, resulting in a loss of BRET and hence allowing quantification of the binding properties of the experimental compounds (**Figure S2**, **B**). The NanoBRET assay represents a more realistic assessment of binding potency in comparison with the ADP-Glo assay, because it simultaneously measures the inhibitors’ ability to penetrate the cells and bind to ULK1/2 in the presence of the cellular environment. Consequently, the results obtained in the NanoBRET assay are a critical step in our testing platform, enabling the selection of compounds for further characterization.

### Structure-activity relationship (SAR) studies & characterization/optimization of ULK1/2 inhibitors

We previously reported the optimization of a trisubstituted pyrimidine scaffold (**Figure 1**) to afford compounds of high potency for ULK1/2 that inhibit autophagy^2^. Our greater goal is to identify compounds with both oral exposure and favorable absorption, distribution, metabolism, and excretion (ADME) properties, while retaining the capacity to inhibit autophagy. To identify novel ULK1/2 inhibitors, we set out to systematically explore a wider chemical space of the R^1^ and R^2^ substituents of the pyrimidine core, utilizing learnings from our prior SAR work and the published co-crystal structures of ULK1/2 ^11, 26^. The ULK2/**SBI-0206965** crystal structure (PDB-ID 6YID) offered an opportunity for optimized spatial conformity within the ATP binding cavity formed in the hinge region between the N- and C-lobes of the kinase domain, as well as possibilities for additional ULK1/2 protein interactions resulting from modifications to the trimethoxy phenyl ring within the solvent-facing region. We also sought to explore the impact of interactions with the hydrophobic back pocket of the ULK1/2 kinase. While designing novel analogs, we also aimed to improve the physical properties of the inhibitors as well as cell penetration and their pharmacokinetic properties. All newly synthesized analogues were evaluated using ADP-Glo and NanoBRET assays, with the results summarized in **Table 1**.

**Figure 1.**
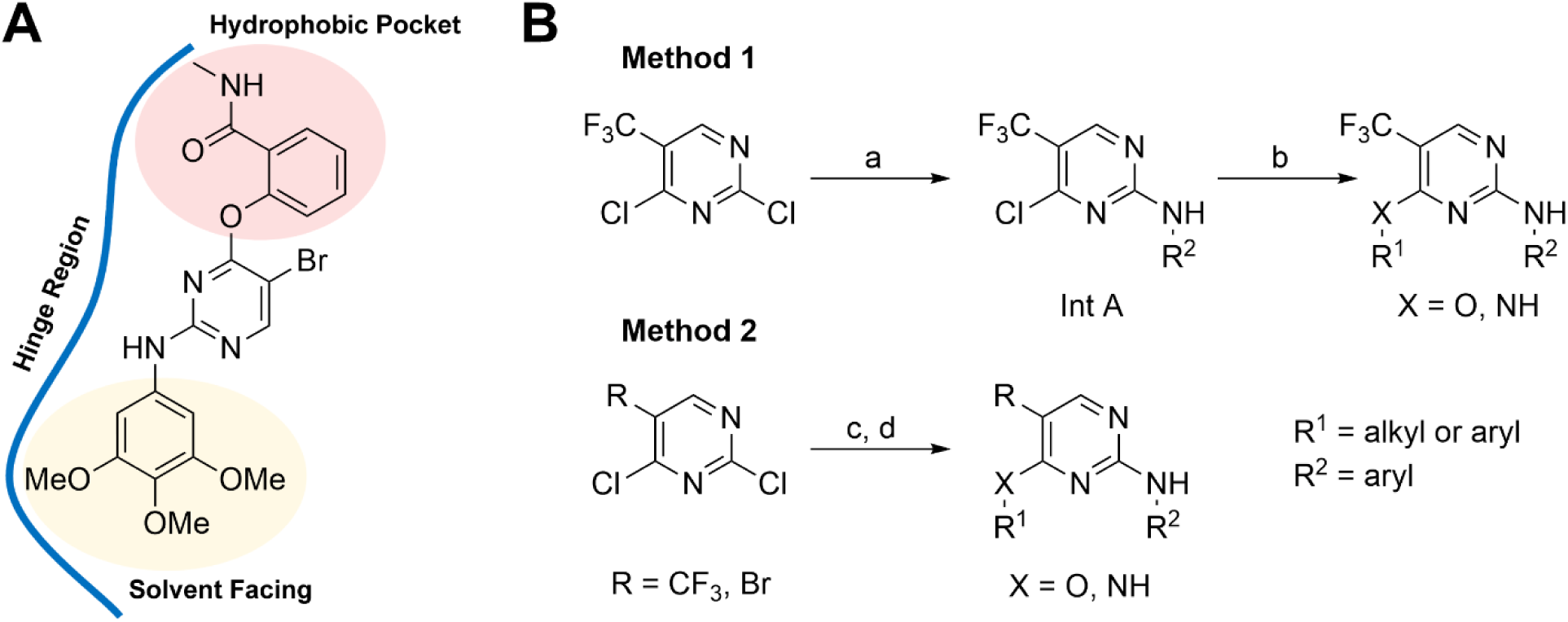
Structure-activity relationship strategy around ULK1/2 fragments. (A) Chemical structural illustration of ULK1/2 small molecule inhibitor **SBI-0206965**. The compound interacts with the ULK1/2 hinge region, occupying the hydrophobic pocket (highlighted in red) and extending to the solvent-facing region (highlighted in yellow). The pyrimidine scaffold of **SBI-0206965** forms hydrogen bonds with the hinge region (blue). (B) General synthetic route for the preparation of ULK 1/2 inhibitors. *Method 1* reagents and conditions: (a) ZnCl_2_ (1.0 M solution in ether), CH_2_Cl_2_-*^t^*BuOH, 0 °C, 1 h; R^2^NH_2_, Et_3_N, 0- rt, 1.5 h (b) R^1^XH, DIPEA, ACN or DMF, µW, 100-130°C, 10-30 min. *Method 2* reagents and conditions: (c) R^1^XH, DIPEA, ACN, µW, 100 °C, 10 min. (d) R^2^NH_2_, AcOH, µW, 120 °C, 10 min.

New amine-containing analogues were synthesized using previously established synthetic routes (**Figure 1B**) ^2, 17^. Earlier studies showed that introducing an amino substituent at the C_4_-position of the pyrimidine scaffold significantly enhanced ULK1 inhibitory activity compared to an ether analogue at the C_4_ position—for example, **SBP-7455** exhibited a 10-fold increase in potency over **SB1-0206965** (IC□□ = 13 nM vs. 130 nM, respectively; **Table 1**). Previously we have described the SAR of an extensive series of ULK1/2 inhibitors where the amine substituent R^2^ has been modified with substituted and unsubstituted aryl, heteroaryl, and bicycloaryl groups. This resulted in compounds that were generally very potent in the ULK1 ADP-Glo assay but exhibited reduced cell potency.^2^ The most potent analog of this series was SBP-7455 with an IC_50_ of 328 nM in the cell-based ULK1 NanoBRET assay. The X-ray structure of ULK2/**SBI-0206965** shows that the R^2^ aryl group occupied the hinge region, sandwiched between the N- and C-lobes, and faced towards solvent. This then offered the opportunity to prepare several more analogs in which the R^2^ group was modified with substituents designed to form productive contacts with nearby polar surface residues such as Asp102 of ULK1 and Asp95 of ULK2. To this end, we investigated a small group of R^2^ substituents and investigated their effects on kinase potency and intracellular activity. The first compound, reference compound **1**, was an analogue of **SBP-7455**, in which the 3,4-dimethoxyphenyl group was replaced with the trimethoxyphenyl substituent, derived from **SBI-0206965** (**Table 1**). **1** demonstrated improved potency, with IC□□ values of 5 nM in the ULK1 ADP-Glo assay and 236 nM in the ULK1 NanoBRET assay. The 6-amino-3,4-dihydroquinolin-2(1*H*)-one derivative **2 (SBP-7501)** had excellent biochemical properties against ULK1 (ADP Glo IC_50_ = 4 nM and ULK1 NanoBRET IC_50_ = 167 nM), while the ring contracted analog **3** was only slightly less potent in both assays. We then investigated a similar scaffold, tert-butyl 6-amino-3,4-dihydroisoquinoline-2(1H)-carboxylate, in both its Boc-protected and deprotected forms. The Boc-protected analogue, **4**, exhibited potent ULK1 kinase inhibition (IC_50_ = 16 nM) but disappointing cell potency (IC_50_ = 8.1 μM NanoBRET). Upon Boc deprotection, the resulting compound **5 (SBP-5147)** showed excellent activity in both the ULK1 ADP-Glo assay (2 nM) and the NanoBRET cellular target engagement assay (47 nM) (**Table 1**). Presumably, the poor cell potency of the Boc-protected analog **4** is a result of its very high lipophilicity (cLogP = 6.28), which reduces its ability to penetrate the cell compared to that of **5** (cLogP = 4.15) (See **Table S1**). The exceptional potency of **5** is likely a result of the tetrahydroisoquinoline nitrogen successfully picking up additional interactions at the protein surface (vide infra).

The CF_3_-group of **SBP-7455** is known to occupy a small, restricted pocket at the back of the hinge region and we decided to reinterrogate this site in case the optimized R^2^ moieties had any effect on how the molecules bound. Consequently, the bromo-analog of compound **1** was prepared, compound **6**, and it was found to be approximately 2-3-fold less active in both assays. Interestingly, the effect of the bromine atom replacement of CF_3_ in compound **2**, was more dramatic in the ULK1 ADP-Glo assay (8-fold reduction in potency compound **6** vs **2**) but showed only a 2-fold potency reduction in the nanoBRET assay. Analog **7** demonstrated a 2-fold improved ULK1 NanoBRET activity relative to **6**. We also synthesized several analogs in which the amine linker (Y = NH in **Table 1**) was replaced with an ether link (Y = O; data not shown). These analogs showed no improvements in the ADPGlo assay and were significantly poorer in the nanoBRET assay.

Removal of the substituent and its replacement with a fused phenyl ring (compound **8**) resulted in a marked loss of potency in both assays. Thus, the trifluoromethyl group at the pyrimidine 5-position appeared to be the preferred substituent and was held constant for the remaining SAR studies.

In the crystal structure of **SBI-0206965** bound to ULK2, it was observed that the *N*-methyl benzamide group did not ideally conform to the ATP binding pocket ^27^. This presented an opportunity to see if alternative R^1^ groups could yield improvements in binding and evaluate the effect of physical properties on cellular activity. To investigate the effect of modifications to R^1^, we elected to keep the R^2^ substituent fixed as the 6-amino-3,4-dihydroquinolin-2(1*H*)-one due to the excellent potency profile of **2**. Firstly, the addition of a methyl group to the 4-N-cyclopropyl substituent afforded the trisubstituted amine analog **9** (Y= -NcPrMe, **Table 1**). Compound **9** had a 3-fold drop in potency vs **2** in the ULK1 ADP-Glo assay but a very significant 30-fold loss of potency in the nanoBRET assay (IC_50_ = 4.9 µM) suggesting the importance of the amine with respect to cellular potency and hydrogen bonding interactions. Removal of the cPr group from **2** yielded **10** which, at an IC_50_ of 144 nM in the ULK1 ADP-Glo assay, was 30-fold lower in potency. Increasing the ring size from cPropyl (**2**, cLogP = 3.85) to cButyl (**11**, cLogP = 4.18) to cPentyl (**12,** cLogP = 4.73) only had a modest effect on the intrinsic inhibitory potency but had a dramatic negative effect on the cellular potency commensurate with the increase in lipophilicity (**Table S1**).

Replacement of the *N*^2^-cyclopropyl moiety of **2** (**SBP-7501),** with ethyl hydrazine, as in **13**, resulted in weaker potency, while *N*^2^-trifluoroethyl substituted analogue **14** resulted in similar ULK1 binding compared to **SBP-7501** (ULK1 ADP-Glo IC_50_ = 6 nM, NanoBRET IC_50_ = 448 nM). Chain extension and incorporation of hydroxy (**15**), cyclopropylmethyl (**16**), 1-cyclopropylmethanone (**17**), or methoxy (**18**) decreased overall potency in relation to **SBP-7501**. Interestingly, a straight chain aliphatic alkylamine substitution as in compound **19** (cLogP = 2.95), led to a compound with favorable biochemical and cellular potency (ULK1 ADP-Glo IC_50_ = 16 nM and NanoBRET IC_50_ = 132 nM; an 8-fold shift). The Boc-protected precursor **20** was also evaluated, showing that the bulky hydrophobic substituent resulted in inferior profiles of both ULK1 ADP-Glo and NanoBRET (IC_50_ = 28 nM and 3259 nM, respectively). In addition, conversion of the amine moiety in **19** to the corresponding cyclopropyl carboxamide derivative (**21**) showed good results in both ULK1 ADP-Glo (IC_50_ = 5 nM) and NanoBRET assays (IC_50_ = 100 nM).

The SAR studies demonstrated that analogs with increased potency can be designed through additional binding interactions with ULK1/2 (e.g. compound **5**). In addition, modulating the lipophilicity of the inhibitors can have a significant effect on the cellular potency (see **Table S1**). Compounds in this series with cLogP >4 often exhibit large potency shifts to lower IC_50_ values in the ULK1 NanoBRET assay relative to their ADP-Glo activity (e.g., compound **4** with a cLogP = 6.28, 500-fold shift; and compound **12** with a cLogP = 4.74, >400-fold shift). In contrast, compounds with cLogP <3.5 generally show smaller shifts between the two assays (e.g., SBI-0206965 with a cLogP = 2.90, 6-fold shift; and compound **10** with a cLogP = 2.44, 16-fold shift).

As indicated in **Table 1**, ULK1 NanoBRET IC_50_ values are right-shifted in comparison with ADP-Glo results. Due to the cellular nature of the NanoBRET assay, factors such as membrane permeability, ATP competition, in-cell protein interactions, and the activation state of the protein can modulate the ability of the compounds to interact with ULK1 ^2^. Based on the ULK1 ADP-Glo and ULK1 NanoBRET IC_50_ values of our previously reported compound **SBP-7455** (13 nM and 328 nM, respectively) ^2^, five compounds in the new series were identified with overall enhanced potency: **1**, **3**, **21**, **SBP-7501**, and **SBP-5147**. Both **19** and **6** were similar to **SBP-7455** in biochemical potency (IC_50_ = 16 nM and 13 nM, respectively). However, these compounds differed in their cellular potency for ULK1, with **19** demonstrating increased binding, and **6** displaying decreased binding to ULK1 as compared to **SBP-7455**. **14** exhibited greater biochemical potency than **SBP-7455** but reduced intracellular target engagement. Both **15** and **7** were less potent in biochemical assays compared to **SBP-7455**. While **15** displayed inferior potency in NanoBRET assays by approximately 2-fold in relation to **SBP-7455** (IC_50_ = 605 nM and 328 nM, respectively), **7** exhibited a similar intracellular profile (**Table 1**).

As the ULK1 homologue ULK2 can compensate for the loss of ULK1 activity ^28^, we subsequently tested our novel inhibitors for activity in ULK2 ADP-Glo and ULK2 NanoBRET assays. As observed with ULK1, ULK2 NanoBRET IC_50_ values were right-shifted compared to ADP-Glo results (**Table S2**). Among the tested compounds, **1**, **21**, **SBP-7501**, and **SBP-5147** were found to be the most effective at both inhibiting ULK2 kinase activity and binding to intracellular ULK2. Moreover, **21** and **SBP-5147** displayed higher binding affinities for ULK2 compared to ULK1, with NanoBRET results for ULK2 improved by roughly 2- and 3-fold, respectively (**Table S2**).

Additionally, selected **Table S2** compounds were tested in the protein thermal shift (PTS) assay to verify their in vitro binding capacity and assess their relative ability to stabilize ULK2, as measured by a shift in protein melting temperatures (Δ*T_m_*). In this assay, a steady increase in temperature (up to 95 °C) unfolds the protein, leading to an increase in fluorescence as the fluorescent dye binds to the exposed hydrophobic sites of ULK2 (**Figure S2**, **C**). Ligands that bind and stabilize the ULK2 protein increase the *T_m_*, and ΔT*_m_* often correlates with the affinity of the ligand ^29, 30^. **SBP-5147** presented the largest thermal shift among the tested compounds (ΔT*_m_* = 14 °C) and also outperformed the previously published compounds, **SBI-0206965** (ΔT*_m_* = 8 °C) and **SBP-7455** (ΔT*_m_* = 9.5 °C) (**Table S3**). **SBP-7501** produced the second largest thermal shift (ΔT*_m_* = 9.5 °C), comparable to that of **SBP-7455**. In conclusion, based on the SAR studies and results from the ADP-Glo, NanoBRET, and PTS assays, **SBP-5147** was identified as the most potent compound in the new series of ULK1/2 inhibitors.

### Structural basis for the potency of SBP-5147

Our SAR analysis identified **SBP-5147** as a potent ULK 1/2 inhibitor. To elucidate its binding mode, we used our established crystallization platform ^11^ and co-crystallized **SBP-5147** with the kinase domains of both ULK1 (16-278) and ULK2 (1-276). The structures were solved at 2.0 and 1.97 Å resolution, respectively. Although the ATP-binding pockets of ULK1 and ULK2 share strong sequence and structural similarity ^11^, the two proteins crystallized in distinct conformations. ULK1 crystallized with two molecules in the asymmetric unit, with a well-structured kinase domain including the activation segment. In contrast, ULK2 crystallized in a dimeric form with two dimers in the asymmetric unit, adopting a conformation reminiscent of the activation loop exchanged state reported for other kinases ^31, 32^. This ULK2 conformation, previously described ^11^, did not affect the ATP site or inhibitor binding mode.

**SBP-5147** engaged the ATP-binding site of both kinases (**Figure 2, A** and **D**), suggesting that it inhibits ULK1 and ULK2 in an ATP-competitive fashion. Key interactions are highlighted in **Figure 2**. The 2-aminopyrimidine core formed bidentate hydrogen bonds with the backbone carbonyl oxygen and amide nitrogen of the hinge cysteine (ULK1 Cys95; ULK2 Cys88). The thiol side chain of this Cys residue was buried within the protein core and thus inaccessible, precluding its use in covalent cysteine-targeted inhibitor design. The *N*-cyclopropyl substituent was rotated out-of-plane from the pyrimidine ring and made hydrophobic contacts with Val30 of the glycine-rich loop. Extending from the scaffold towards the solvent-exposed surface of the binding cleft, the protonated 1,2,3,4-tetrahydroisoquinoline forms an ionic interaction with the side chain of a conserved Asp residue (ULK1 Asp102; ULK2 Asp95). This feature distinguishes **SBP-5147** from our previous ULK1/2 chemical probe, **SBI-0206965**, whose trimethoxy phenyl ring could not participate in similar contacts. Docking studies with SBP-7501, which replaces the 1,2,3,4-tetrahydroisoquinoline with a 3,4-dihydroquinolinone moiety and is 4-fold less potent than SBP-5147 in the ULK1 NanoBRET assay, suggested that the 3,4-dihydroquinolinone cannot directly engage the Asp side chain in ULK1/2, potentially accounting for SBP-7501’s reduced potency (**Figure S3**). Finally, the trifluoromethyl group at pyrimidine position 5 was placed against the gatekeeper methionine in the back pocket (ULK1 Met92; ULK2 Met85), making favorable hydrophobic interactions. Overall, these structural studies establish the basis for SBP-5147’s potency against ULK1/2, define its unique interaction profile, and highlight opportunities to further exploit hinge and pocket residues in future inhibitor design.

**Figure 2.**
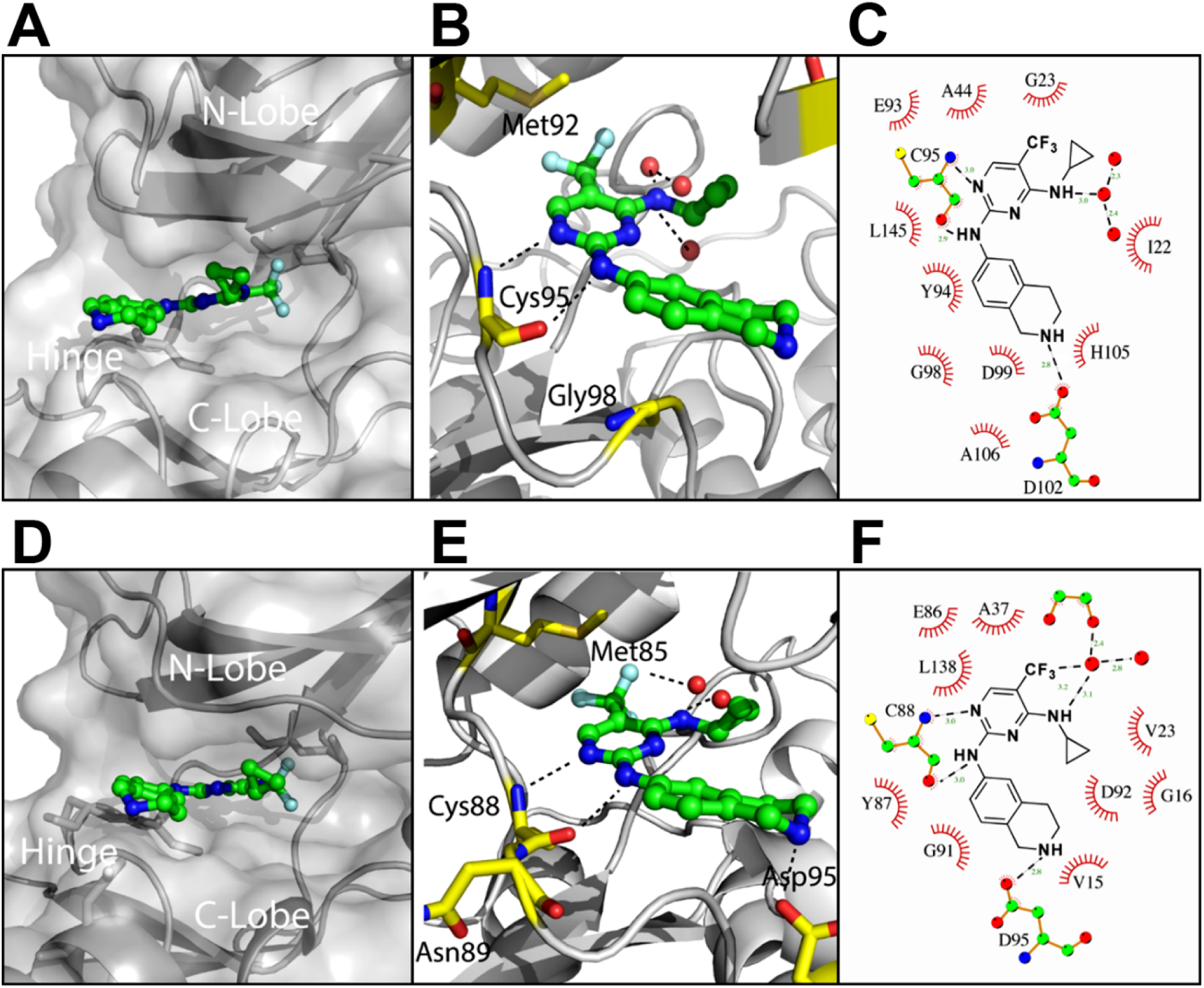
The binding of SBP-5147 to the kinase domain of ULK1 and ULK2 shares a common inhibition mode. (A) Crystal structure of **SBP-5147** bound to the ATP-binding site of ULK1. (B) **SBP-5147** key interactions with the ULK1 kinase domain. (C) Schematic of the interaction profiles of **SBP-5147** to the ULK1 kinase domain. (D) Crystal structure of **SBP-5147** bound to the ATP-binding site of the ULK2 kinase domain. (E) **SBP-5147** key interactions with the ULK2 kinase domain. Polar contact counterparts to those observed with ULK1 are retained in the structure of the kinase domain of ULK2. (F) Schematic of the interaction profiles of **SBP-5147** to the ULK2 kinase domain.

### Evaluation of downstream target engagement by ULK1/2 inhibitors in HEK293T cells

We next evaluated the ability of the top ten most potent compounds to inhibit autophagy-activating proteins downstream of the ULK1/2 preinitiation complex by assessing two substrates of the ULK1/2 kinase, Beclin-1 and VPS34. Upon autophagic stimuli, phosphorylation of Beclin-1 and activation of the pro-autophagy VPS34 complex by ULK1/2 are required for adequate autophagy induction in mammals ^33^. Hence, we quantified the inhibition of Beclin-1 phosphorylation at Ser15 and VPS34 phosphorylation at Ser249 in HEK293T cells expressing Myc-tagged kinase inactive (KI) or wildtype (WT) ULK1, and either Flag-tagged Beclin-1 or VPS34. HEK293T cells were selected due to their high transfection efficiency.

Transfected cells were treated with either DMSO or ULK1/2 inhibitors at 10 µM for 1 h and the phosphorylation states of Beclin-1 at S15 and VPS34 at S249 were assessed via Western blot (**Figure 3**, **upper panels**). Cell lysates were immunoblotted with antibodies to phospho- and total Beclin-1 or phospho- and total Vps34, in addition to Myc-tag antibodies, which verify ULK1 expression. The bands representing total and phosphorylated Beclin-1 and VPS34 proteins were quantified by densitometry and normalized to DMSO-treated cells (**Figure 3**, **bottom panels**). Downstream target engagement was observed for all evaluated compounds as measured by a decrease in the ratio of phospho- to total Beclin-1 or phospho- to total Vps34. However, compounds **SBP-5147**, **SBP-7501,** and **1** were the most effective, inhibiting phosphorylation of the target proteins by roughly 80%, 70%, and 60%, respectively. In comparison, our first-generation ULK1 inhibitor, **SBI-0206965** ^27^, which served as a positive control, reduced the phosphorylation of Beclin-1 and Vps34 by only 27% and 45%, respectively. Together, these data indicate that **SBP-5147** and **SBP-7501** display enhanced ULK1 kinase inhibition as compared to our previous molecules. (**Figure 3**, **lower panels**).

**Figure 3.**
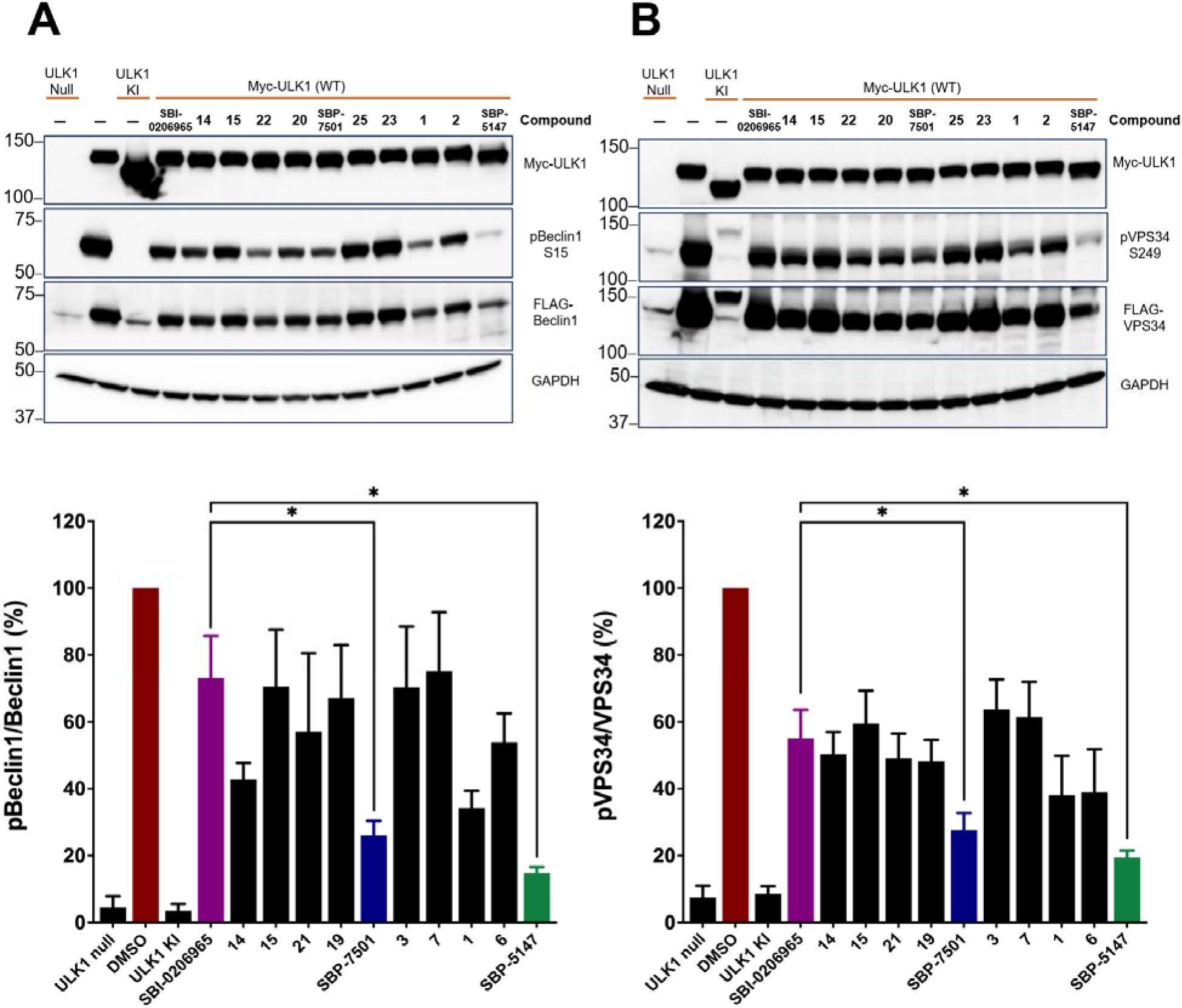
ULK1/2 inhibitors decrease the phosphorylation of downstream autophagy target proteins. Western blot analyses (upper panels) and quantification (lower panels) of Beclin1 and Vps34 phosphorylation upon treatment with ULK1/2 inhibitors. HEK293T cells were transfected with Myc-tagged kinase inactive (KI) or wildtype (WT) ULK1 and WT Flag-tagged (**A**) Beclin1 or (**B**) Flag-tagged Vps34 for 24 h. Subsequently, cells were treated with DMSO or the indicated ULK1/2 inhibitors at 10 µM for 1 h. Cell lysates were immunoblotted with the indicated antibodies. The density of each band was quantified for DMSO and compound-treated samples. Data are expressed as the ratio of p-Beclin1/Beclin1 or p-VPS34/VPS34 normalized to GAPDH and DMSO-treated samples. Data represent the mean ± SEM of three independent experiments. *P<0.05 vs **SBI-0206965** by unpaired t-test with Welch’s correction.

Overall, these findings confirm that our novel ULK1/2 inhibitors engage the intended intracellular targets and effectively inhibit phosphorylation of downstream substrates of ULK1/2. Moreover, according to the results of the ULK1/2 ADP-Glo and NanoBRET assays, ULK2 PTS assay, and the observed decrease in phosphorylation of Beclin-1/VPS34, we concluded that **SBP-5147** and **SBP-7501** are the most potent ULK1/2 inhibitors in our new series and thus were selected for further evaluation and characterization.

### ULK1/2 inhibitors decrease autophagic flux in A549 cells

Autophagic flux is a measurement of the rate of autophagic degradation. To investigate autophagic flux in cells and therefore demonstrate autophagy inhibition by **SBP-5147** and **SBP-7501**, we employed a flow cytometry-based assay using A549 cells expressing a tandem-labeled mCherry-GFP-LC3 chimeric reporter protein to quantify the ratio of autophagosomes to lysosomes (**Figure 4, A**) ^34^. A549 cells were selected because they are easy to grow (doubling time is roughly 20 h) and are a well-characterized NSCLC cell line.

**Figure 4.**
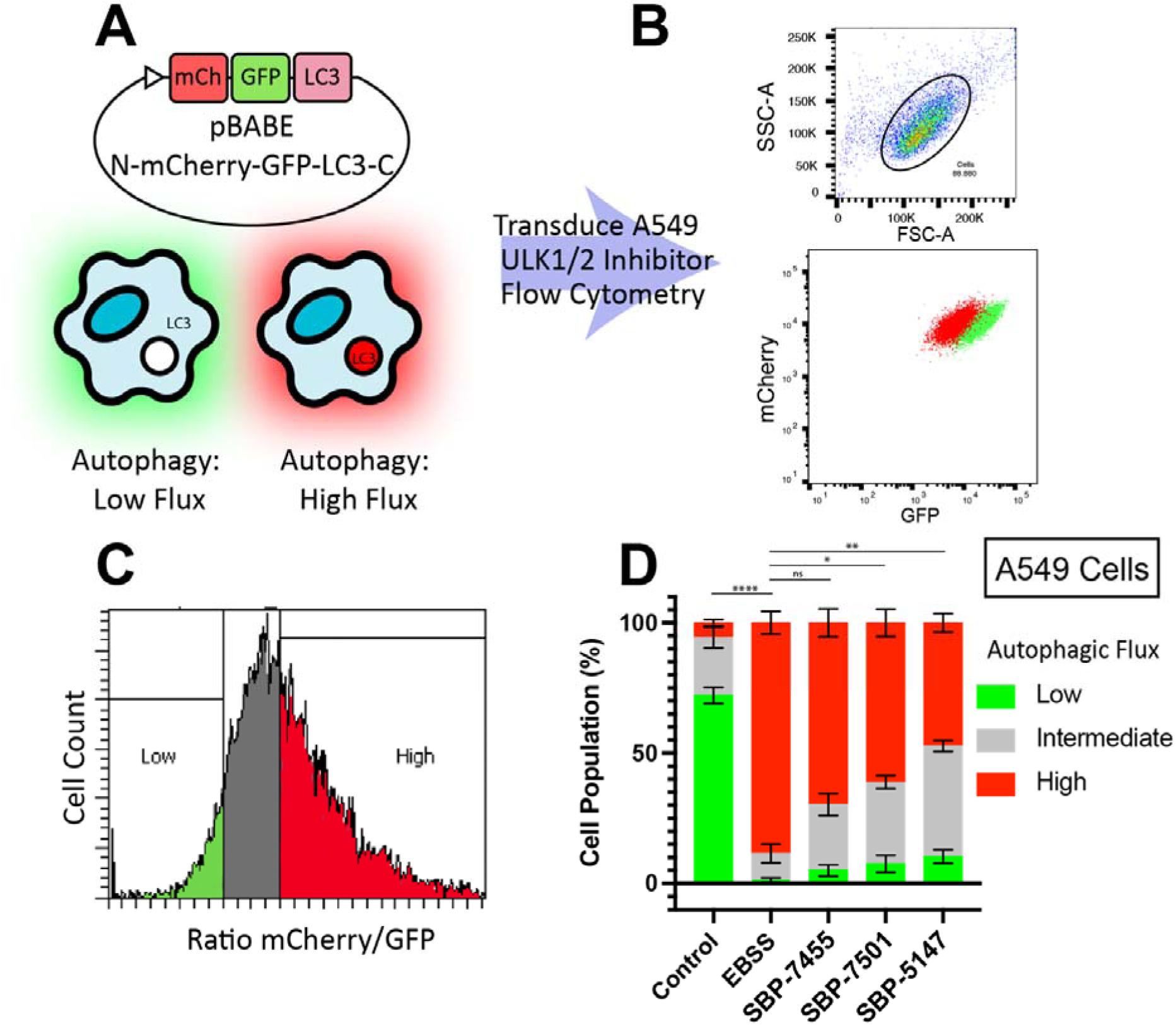
ULK1/2 inhibitors decrease autophagic flux in A549 cells. (**A**) A549 cells expressing the reporter pBABE-mCherry-EGFP-LC3B were used to evaluate autophagic flux. GFP exhibits increased sensitivity to the acidic environment of the autolysosome compared to mCherry. Under conditions of high autophagic flux, fusion of autophagosomes with the lysosome increases the mCherry/GFP ratio in the cell. (**B**) A549 cells were incubated in growth media (control), starvation media (EBSS), or starvation media plus ULK1/2 inhibitors at 10 μM for 18 h prior to flow cytometry analysis. The relative LC3 distribution was used as a measure of autophagic flux in the cells: low flux (green) cells under growth media compared to high flux (red). (**C**) Representative histogram of A549 cells treated with ULK1/2 inhibitors, showing populations of low (green), intermediate (grey), and high (red) autophagic flux. (**D**) Analysis of three independent experiments, data indicate the mean ± SEM, ****P≤0.0001, ns= P≥0.05, *P≤0.05, **P≤0.01 by two-tailed paired t-test, high autophagic flux population vs control.

Since the GFP tag is sensitive to the acidic environment of the lysosome (pH 5), GFP fluorescence is extinguished upon autophagosome-lysosome fusion, while the mCherry tag is acid-insensitive and stable until it is degraded (**Figure 4, B**). Therefore, under nutrient-stressed conditions, fusion of autophagosomes with lysosomes increases the mCherry-GFP ratio in the cell (high autophagic flux). Accordingly, autophagy inhibition is measured by a decrease in the cellular ratio of mCherry to GFP fluorescence (**Figure 4, C**). In this assay, A549 mCherry-GFP-LC3 cells were maintained in enriched media (control), starvation media (EBSS), or starvation media plus ULK1/2 inhibitors at 10 µM for 18 h. Wildtype A549 cells were used as non-fluorescent control. In the absence of autophagy stimuli, approximately 60% of transduced A549 cells displayed low autophagic flux characterized by a low ratio of mCherry to GFP. On the other hand, after incubation in starvation media, roughly 90% of cells presented high autophagic flux (high ratio of mCherry to GFP). Treatment of cells in starvation media with **SBP-5147**, **SBP-7501**, and **SBP-7455**, resulted in a significant decrease in the mCherry to GFP ratio (i.e., autophagic flux), confirming autophagy inhibition by our ULK1/2 inhibitors (**Figure 4, D**).

### ULK1/2 inhibitors reduce the viability of NSCLC cells but are not cytotoxic against non-tumorigenic lung cells

Subsequently, we examined the cytotoxic activity of **SBP-5147** and **SBP-7501** as single agents to prevent the growth of NSCLC cells. For our viability studies, in addition to A549 cells (*KRAS G12S/STK11*-mutant), we selected lung cancer cell lines that present clinically relevant oncogenic alterations ^35^. Therefore, H1373 (*KRAS G12C*), HCC827 (*EGFR* exon 19 deletion), and H1975 (*EGFR L858R/T790M, KRAS G12D*) were used in this assay to determine if cellular mutational status could predict susceptibility to ULK1/2 inhibition. Moreover, we assessed the viability of BEAS-2B, a non-tumorigenic epithelial lung cell line, following treatment with our ULK1/2 inhibitors. BEAS-2B is commonly utilized as an in vitro cellular model for evaluating chemicals and biological agents with potential pulmonary toxicity or lung carcinogenicity ^36^.

Cells were seeded in 384-well plates spotted with 3-fold serial dilutions of the test compounds, DMSO, or Staurosporine, (a non-selective protein kinase inhibitor used as a positive control) and incubated for 72 h at standard cell culture conditions. Cell viability was assessed using the ATP-depletion assay, CellTiter-Glo^®^. For each compound, the cell viability compared to DMSO controls was analyzed and IC_50_ values were determined for each cell line. The IC_50_ values of **SBP-5147** and **SBP-7501** in each lung cancer cell line are depicted in **Figure 5** and range between 42 to 161 nM. Conversely, BEAS-2B cells were not particularly susceptible to ULK1/2 inhibition, with only the highest concentrations of **SBP-5147** (30 and 10 µM) and **SBP-7501** (30 µM) reducing cell viability by ≥ 50% (**Figure 5**, dotted line and dotted circles). Our findings indicate that **SBP-5147** and **SBP-7501** potently kill NSCLC cells in culture regardless of mutational status but only reduce the viability of non-tumorigenic epithelial lung cells at high concentrations.

**Figure 5.**
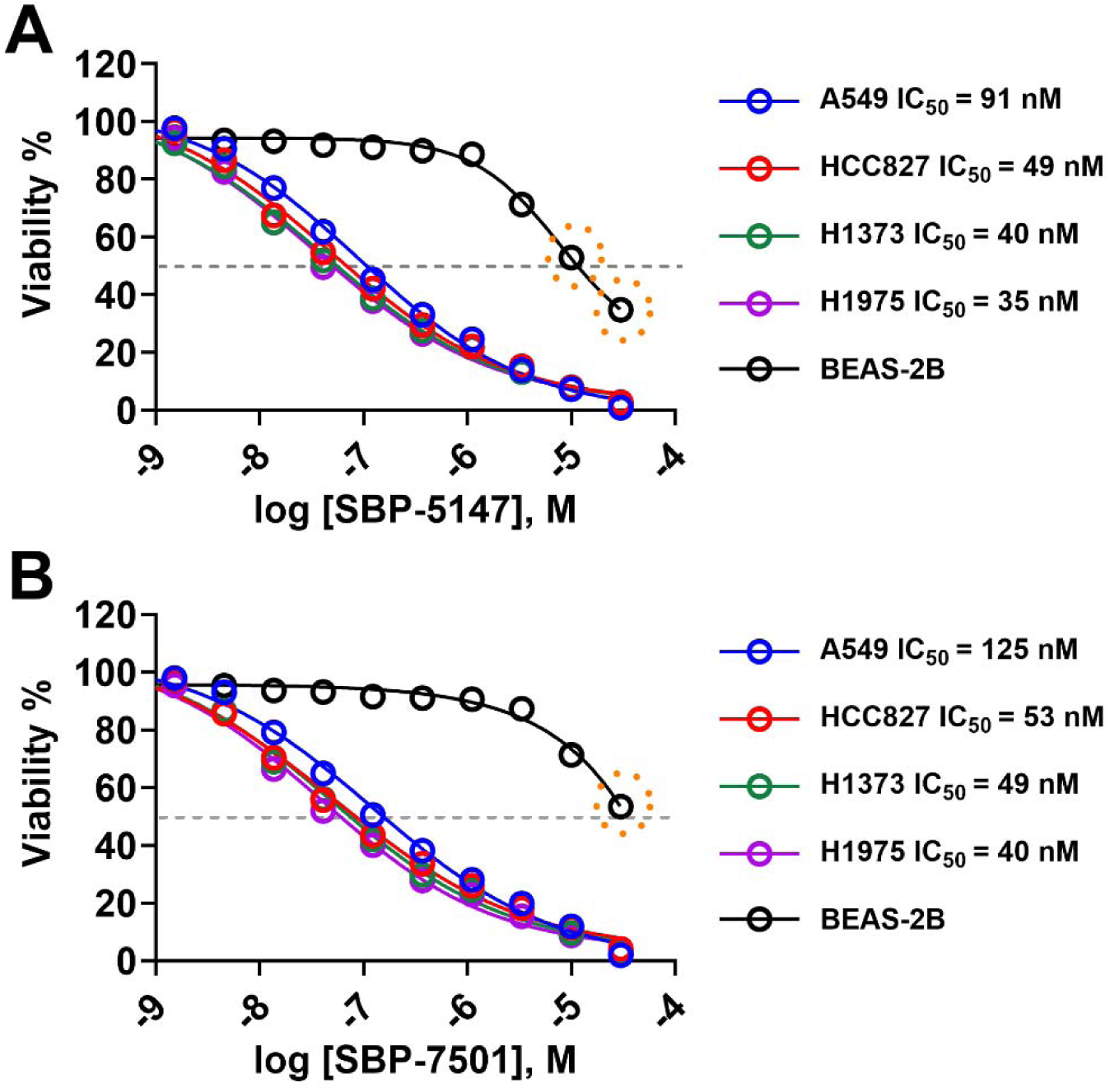
ULK1/2 inhibitors reduce the viability of NSCLC cells as single agents but do not exhibit the same cytotoxicity to non-tumorigenic lung cells (BEAS-2B). Cells were incubated for 72 h at standard cell culture conditions in assay plates spotted with test compounds **SBP-5147** (**A**) and **SBP-7501** (**B**) serially diluted 1:3 from 30 µM to 1.5 nM or vehicle control (DMSO). Cell viability was assessed by the ATP-depletion assay CellTiter-Glo®. The percentages compared to DMSO vehicle control were curve-fitted using nonlinear regression (log [inhibitor] vs. response, variable slope, four parameters). The normalized dose-response curves indicate the mean ± SEM of three independent experiments performed in quadruplicate.

### Pharmacokinetic (PK) analysis of SBP-5147 & SBP-7501

To evaluate the PK properties and oral availability of **SBP-5147** and **SBP-7501**, we administered a single dose (10 mg/kg) via oral gavage to mice and assessed plasma exposure over 24 h by LC-MS/MS. The PK analysis of **SBP-5147** (**Figure 6, A**) showed that the time to peak plasma concentration (T_max_) was 15 min and the maximum plasma concentration (C_max_) was 664 nM, a value approximately 14-fold higher than its NanoBRET IC_50_ value for ULK1 (IC_50_ = 47 nM) and 39-fold higher than its NanoBRET IC_50_ value for ULK2 (IC_50_ = 17 nM). Moreover, the plasma concentration of **SBP-5147** remained above the ULK1/2 NanoBRET IC_50_ value for at least 8 h post-dosing. These data indicate that the plasma levels of **SBP-5147** are sufficient for in vivo target engagement. Additionally, **SBP-7501** (**Figure 6, B**) demonstrated a T_max_ at 30 min and C_max_ of 1440 nM, a value roughly 9-fold greater than its NanoBRET IC_50_ value for ULK1 (IC_50_ = 167 nM) and 8-fold greater than its NanoBRET IC_50_ value for ULK2 (IC_50_ = 174 nM). The plasma concentration of **SBP-7501** persisted above the ULK1/2 NanoBRET IC_50_ for approximately 4 h post-dosing. Compared to **SBP-7455** which had a C_max_ of 990 nM and T_max_ of 1 h when dosed at 30 mg/kg orally ^2^, both compounds presented remarkably improved PK properties even when dosed at a lower dose (10 mg/kg vs. 30 mg/kg). **SBP-5147** was selected for further pharmacodynamics (PD) analysis as it demonstrated that the 8 h trough levels exceeded its ULK1/2 NanoBRET IC_50_ values following 10 mpk oral dosing in mice.

**Figure 6.**
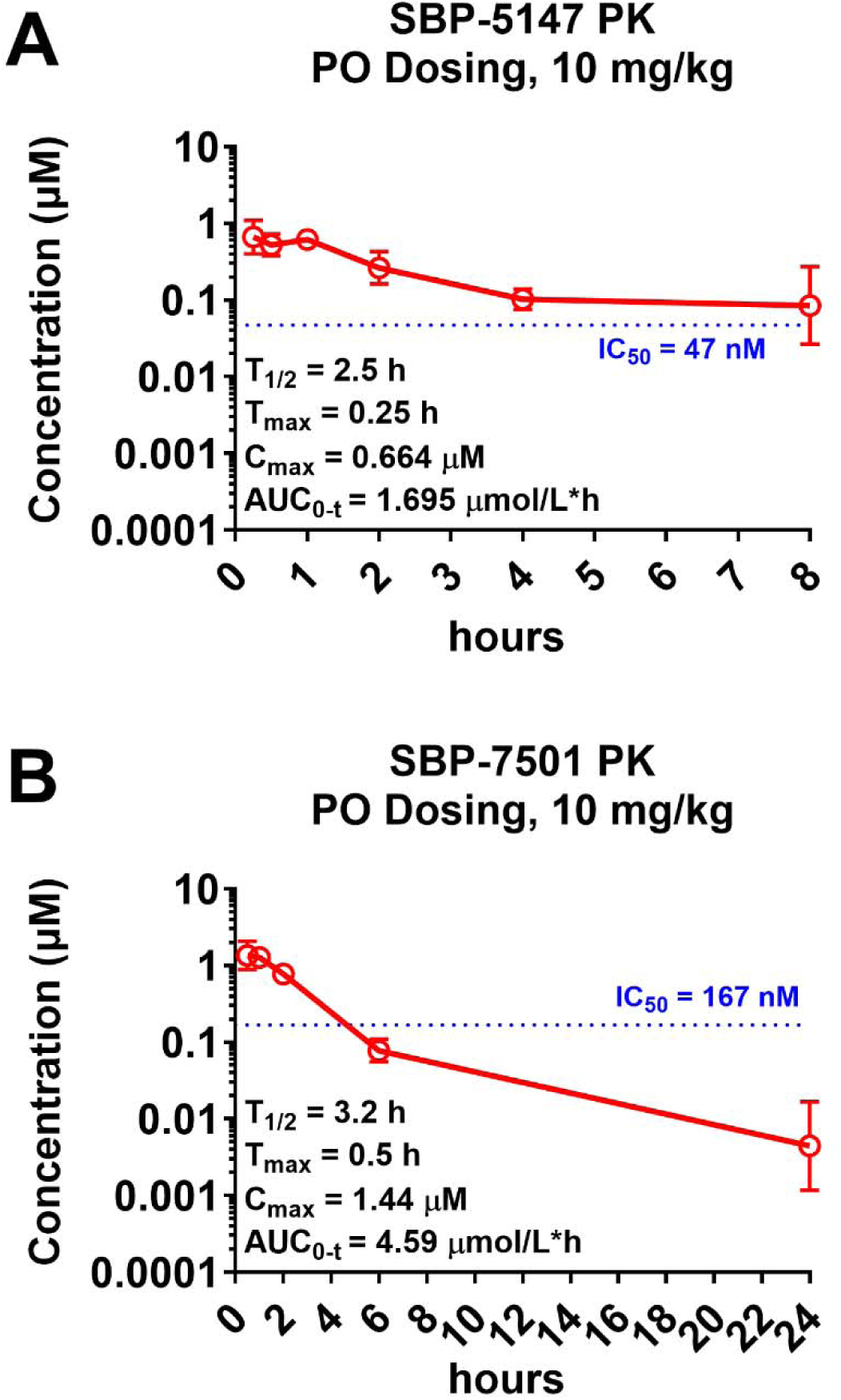
PK analyses of (A) SBP-5147 & (B) SBP-7501. Mice were dosed orally with the indicated compounds (10 mg/kg). Blood samples were collected retro-orbitally and plasma was separated by centrifugation at the indicated time points post-dosing. The compound concentration in the plasma was determined by LC-MS/MS analysis and data were analyzed using PKSolver software. Data represent the geo mean ± geo SD of N=3, log10 scale. The IC_50_ values of **SBP-5147** and **SBP-7501** in the ULK1 NanoBRET target engagement assay are indicated at the dotted lines.

### In vivo target engagement of SBI-5147

ATG13 is a key subunit of the ULK1 preinitiation complex and is essential for autophagy initiation ^37^, serving as an adaptor protein to recruit ULK1, FIP200, and ATG101 for autophagy induction ^38^. Moreover, studies have shown that in addition to primary inhibition of ULK1 kinase activity, suppressing ATG13 binding to FIP200, ULK1, or ATG101 disassembles the ULK1 complex ^38^. ATG13 and ATG101 are known substrates of ULK1 and in previous reports ^27^ we have characterized the loss of both ATG13 phosphorylation and total ATG13 protein levels in cells following treatment with the ULK1 inhibitor, **SBI-0206965** ^27^. Therefore, disruption of the ATG13-ATG101 interaction impairs autophagy initiation and progression ^37–39^.

Herein we demonstrated that **SBP-5147** effectively inhibits phosphorylation of downstream substrates of ULK1 kinase activity (Beclin-1 and Vps34) and reduces autophagy flux in nutrient-challenged A549 cells. Therefore, based on the favorable in vitro/PK profiles of **SBP-5147** and the role of ATG13-ATG101 interaction in autophagy induction, we investigated the effects of this compound on ATG13 and ATG101 protein levels in tissues of mice treated with a single dose of **SBP-5147** as a readout of autophagy inhibition in vivo and a PD biomarker for ULK1 activity. Mice were dosed via oral gavage (10 mg/kg) or vehicle and sacrificed at 2, 4, 8, and 24 h post-dosing. Liver and lung samples were flash frozen in liquid nitrogen and in vivo target engagement was assessed by immunoblotting for ATG13 and ATG101.

Our in vivo findings showed a progressive and significant decrease in ATG13 and ATG101 protein expression at 2 (data not shown), 4-, 8-, and 24-h post-dosing, both in the liver and the lung (**Figure 7, A** & **B**). The specific loss of both ATG13 and ATG101 following drug treatment demonstrates on-target activity of **SBP-5147** and represents a PD readout of this ULK1/2 inhibitor. At 24 h post-dosing, **SBP-5147** has been almost eliminated from plasma (**Figure 6, A**), however, ATG13 and ATG101 levels are still significantly reduced in the liver and lung compared to vehicle-treated samples, indicating that the effects of **SBP-5147** persist following plasma clearance. As the ATG13-ATG101 interaction is required for autophagy ^38^, the loss of these proteins abolishes autophagic flux in vivo. Consequently, our results show that this novel compound has the potential to successfully modulate autophagy both in vitro and in vivo.

**Figure 7.**
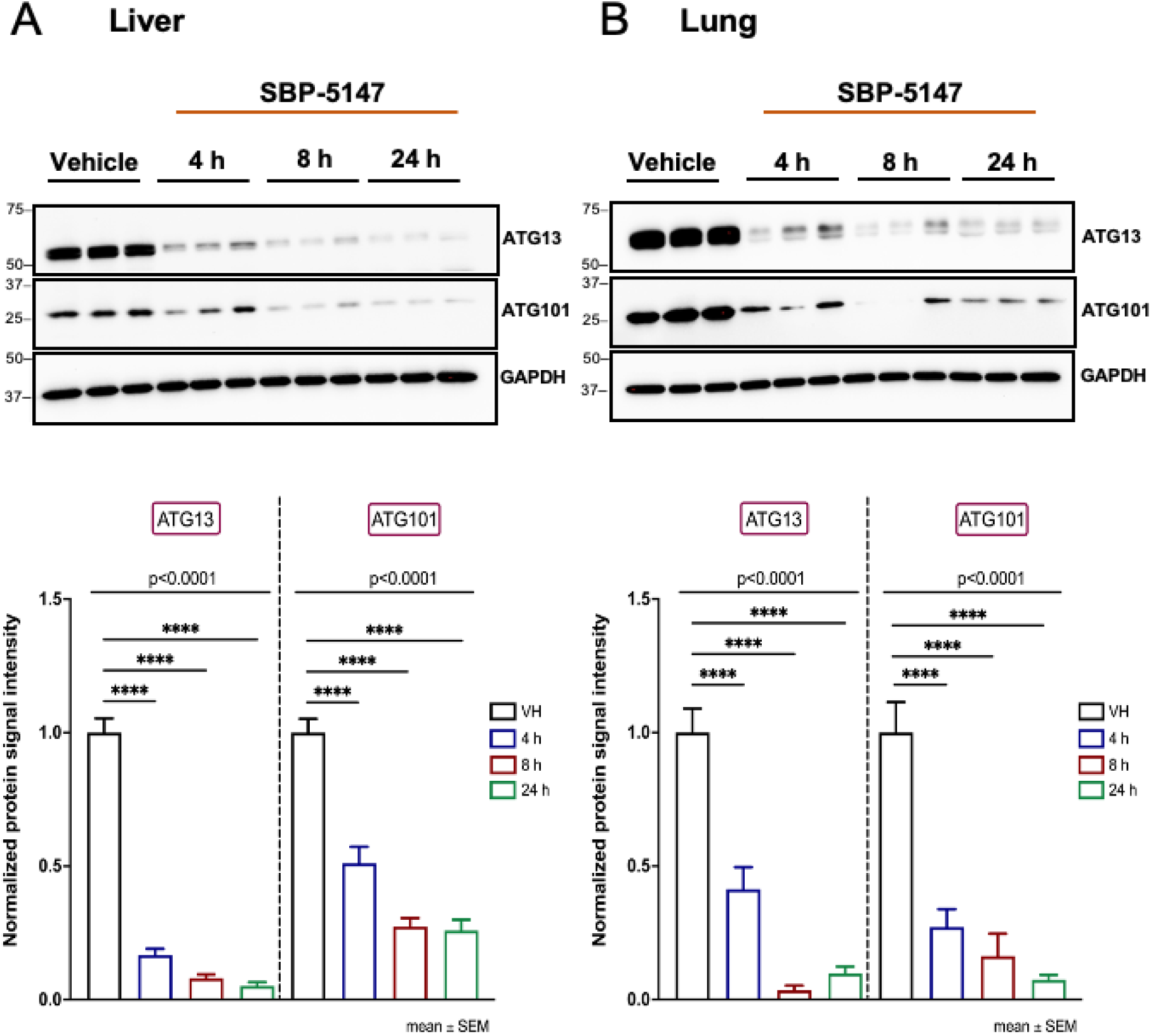
PD analyses of SBP-5147. C57BL/6 mice were dosed orally with **SBP-5147** (10 mg/kg) or vehicle. Tissue samples were harvested at the indicated time points. Tissue lysates were used to evaluate in vivo target engagement of **SBP-5147** by immunoblotting ATG13 and ATG101 in the (**A**) liver and (**B**) lung (upper panels). Lower panels indicate their respective densitometry analyses. The density of protein signals was quantified for vehicle and compound-treated tissue samples and normalized to GAPDH and vehicle-treated samples. A representative image of three independent immunoblotting experiments is shown. Data represent the mean ± SEM of N=3. ****P≤0.0001 by ordinary one-way ANOVA. Stars above horizontal bars: ****P≤0.0001 by Dunnett’s multiple comparisons analyses vs vehicle (post hoc).

### ULK1/2 inhibitors induce apoptosis in A549 & H1975 cells

As this series of ULK1/2 inhibitors effectively kills NSCLC cells in vitro, we investigated the mechanisms of cell death induced by **SBP-5147** and **SBP-7501** in A549 and H1975 cells. The kinetics of cell death induced by **SBP-5147** and **SBP-7501** were analyzed using a live-cell imaging and DNA-staining strategy in which a dye, Sytox Green, is excluded from healthy cells. Dying cells become permeable to this dye upon cell membrane disruption ^40^.

In this assay, cells are treated with inhibitors of specific cell death pathways as well as our ULK1/2 inhibitors. If cell death is attenuated, we can conclude that the pathway that was inhibited contributes to ULK1/2-mediated cell death. We selected concentrations of **SBP-5147** and **SBP-7501** that would induce > 60% of cell death in 24 h in A549 and H1975 cells (**Figure 8, A** and **Figure S4**, **A**). Cells were co-treated with our ULK1/2 compounds as well as a series of inhibitors that target caspases or other signaling proteins of regulated cell death (**Figure 8, A** and **Figure S4**, **A**). To assess apoptosis, we used Emricasan and zVAD-fmk ^41–43^ (pan-caspase inhibitors). To examine necroptosis, we treated the cells with Necrostatin-1 (RIPK1 inhibitor) ^44, 45^, and Necrosulfonamide (MLKL inhibitor) ^46^. Finally, we utilized VX-765, which targets primarily inflammatory caspases ^47^, and an NLRP3-inflammasome inhibitor, MCC950 ^48^ to evaluate pyroptosis.

**Figure 8.**
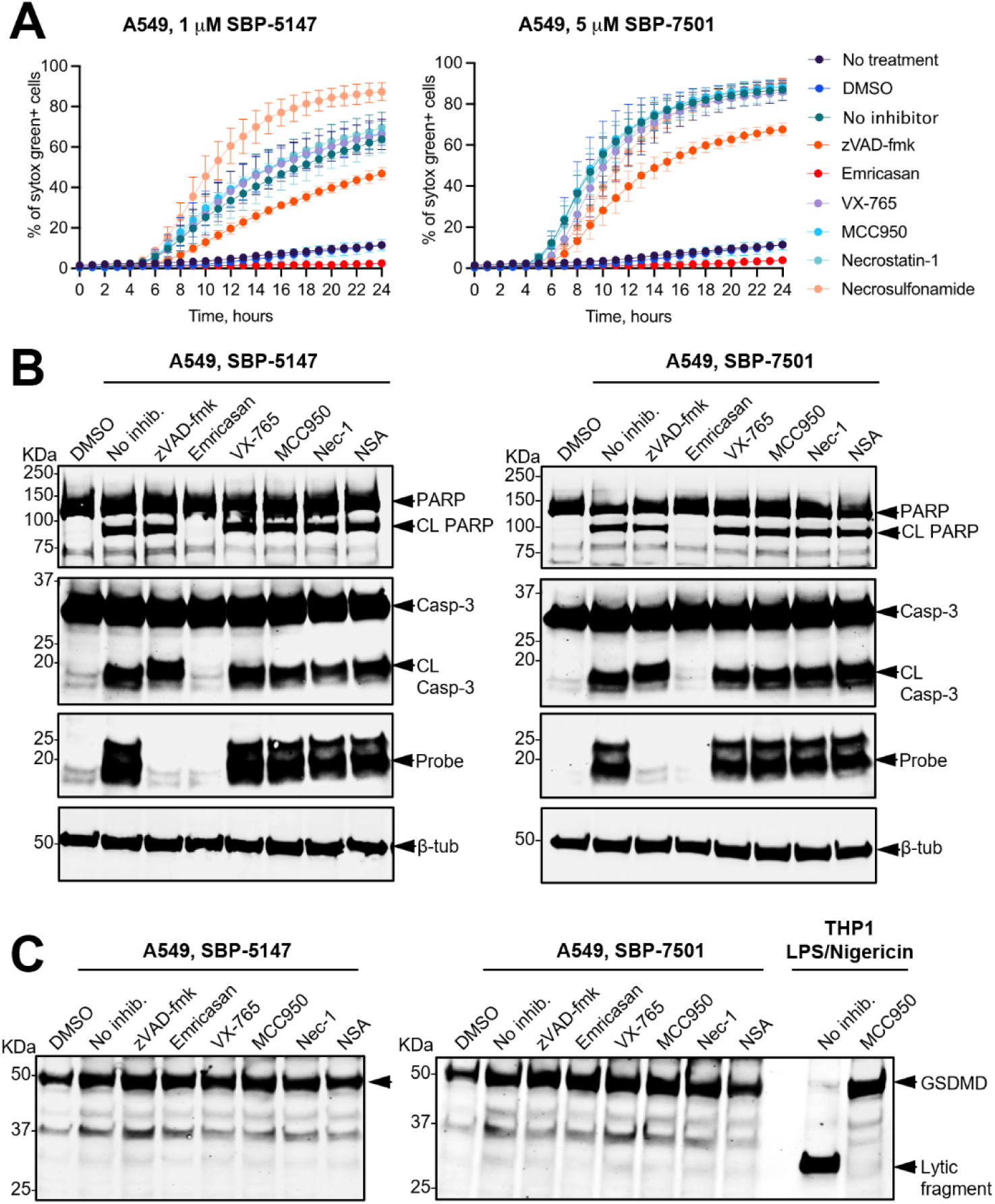
ULK1/2 inhibitors induce apoptosis in A549 cells. **(A)** Cell death assay. A549 cells were treated for 1 h with inhibitors that target caspases or other cell death signaling proteins (zVAD-fmk, Emricasan, MCC950, Necrosulfonamide, VX-765, and Necrostatin-1). Subsequently, cells were treated with **SBP-5147**, **SBP-7501**, or DMSO and imaged for 24 h. Total cell numbers were calculated by Hoechst staining and death cells were detected by sytox green staining. The graph shows the mean ± SD of three independent experiments. (**B**) Apoptosis protein markers were detected by immunoblotting. Cells were treated as described above (panel A) and lysates were collected after 8 h. (**C**) Detection of the pyroptosis effector protein, gasdermin D. THP-1 cells were treated with LPS for 4 h followed by Nigericin for 2 h, with or without MCC950. Western blots are representative of two independent experiments. The probe biotin-ahx-DEVD-AOMK, which reveals active caspase and cleaved proteins, is identified by “CL”.

In A549 cells, treatment with Emricasan completely abrogated cell death and zVAD-fmk treatment showed partial inhibition (**Figure 8, A**). Both caspase inhibitors blocked ULK1/2 inhibitor-induced cell death in H1975 cells, while VX-765 showed limited inhibition (**Figure S4**, **A**). These results indicate that **SBP-5147** and **SBP-7501** induce caspase-dependent apoptotic cell death in both cell lines. Moreover, compound-induced cell death was not affected by compounds targeting necroptosis or the inflammasome. These data indicate that ULK1/2 inhibitors induce cell death in NSCLC cells through an apoptotic mechanism that excludes necroptosis or pyroptosis.

In an orthogonal assay, we examined the activation of the apoptotic pathway using Western blots to assess the cleavage of two proteins associated with programmed cell death, poly ADP-ribose polymerase (PARP) and caspase 3 (**Figure 8, B** and **Figure S4**, **B**). We observed that treatment with **SBP-5147** and **SBP-7501** induced PARP and caspase 3 cleavage, which could be blocked by treatment with apoptosis inhibitors. Overall, these data mirrored the results of the Sytox Green cell death assay and demonstrated an apoptotic cell death mechanism for **SBP-5147** and **SBP-7501**.

Finally, because caspase inhibitors can have non-specific effects on pyroptotic caspases, we sought to determine if the cell death mechanisms induced by **SBP-5147** and **SBP-7501** included a pyroptotic component as well. Pyroptosis is a cell death mechanism regulated by inflammatory caspases through cleavage of the cell death effector protein, gasdermin D (GSDMD), to release its lytic fragment ^49, 50^. Since pyroptosis is triggered by inflammation and thus is usually seen in inflammatory cells such as macrophages, THP-1 cells are typically used to study pyroptotic cell death due to their monocytic and macrophage-like characteristics ^51, 52^. Therefore, THP-1 cell lysates were utilized as positive control for GSDMD cleavage. Pyroptosis was induced in these cells via treatment with lipopolysaccharide for 4 h, followed by Nigericin for 2 h, with or without MCC950. While the GSDMD lytic fragment was observed in THP-1 cells undergoing pyroptosis, we did not identify this marker in A549 or H1975 cells when treated with ULK1/2 compounds (**Figure 8, C** and **Figure S4**, **C**). In addition, although VX-765 attenuated cell death in H1975, the lack of cleaved GSDMD suggests an off-target effect on apoptotic caspases rather than a pyroptotic mechanism.

In summary, our findings indicate that **SBP-5147** and **SBP-7501** induce cell death in NSCLC cells by archetypal apoptosis instead of activating inflammatory-induced cell death pathways. In lung cancer, therapy-induced cancer cell death generates an inflammatory microenvironment that promotes immune escape, tumor angiogenesis, tumorigenesis, and other mechanisms that contribute to tumor progression, metastasis, and development of resistance to subsequent therapies ^53^. Consequently, from a tumor microenvironment standpoint, our data suggest that our chemical probes may present therapeutic utility for adjuvant treatment of NSCLC.

### NSCLC cells treated with SBP-5147 present increased levels of total MHC class I (HLA-A, -B, -C)

Loss of MHC-I antigen presentation machinery (APM) is a common occurrence in cancer, and it is associated with impaired responses to immunotherapy and ultimately worse clinical outcomes ^54^. A study by Deng et al. on NSCLC has demonstrated a mechanism of immune evasion in *LKB1*-mutant lung tumors whereby the expression of components of the immunoproteasome machinery that processes neoantigens is suppressed, reducing the cell-surface expression of MHC-I molecules ^22^. Furthermore, the authors showed that co-treatment with ULK1 inhibitors and immune checkpoint inhibitors (ICIs) results in tumor regression and restores anti-tumor immunity. In pancreatic cancer, it has been shown that MHC-I molecules are selectively targeted for lysosomal degradation by an autophagy-dependent mechanism, whereas inhibition of autophagy restores cell surface MHC-I APM, improves anti-tumor responses, and reduces tumor growth ^55^.

As these data support a therapeutic mechanism where autophagy inhibition sensitizes cancer cells to immunotherapy, we examined the effects of treatment with our top compound, **SBP-5147**, on MHC-I expression in NSCLC cells. NSCLC cells were treated with either DMSO or **SBP-5147** at 15 nM or 30 nM for 72 h in standard cell culture conditions. We selected the concentrations of **SBP-5147** based on the IC_50_ values determined in our ULK2 NanoBRET and cell viability studies and utilized the same NSCLC cell lines, H1373, HCC827, A549, and H1975. Subsequently, cellular lysates were isolated and immunoblotted with an antibody to HLA Class 1 ABC.

H1373, HCC827, and A549 cells displayed similar baseline levels of total MHC-I protein, which were almost undetectable via Western blot (**Figure 9, A**). However, treatment with **SBP-5147** at 30 nM significantly increased total MHC-I protein expression across the three cell lines, with H1373 and HCC827 presenting the most remarkable changes in comparison with A549 cells (**Figure 9, A** and **B**). The same effect was observed in H1373 and HCC827 cells upon treatment with 15 nM of **SBP-5147** (**Figure 9, A** and **B**). Compared to the other cell lines, H1975 presented higher baseline levels of total MHC-I protein, and **SBP-5147** treatment had limited effects on MHC-I upregulation (**Figure 9, A** and **B**).

**Figure 9.**
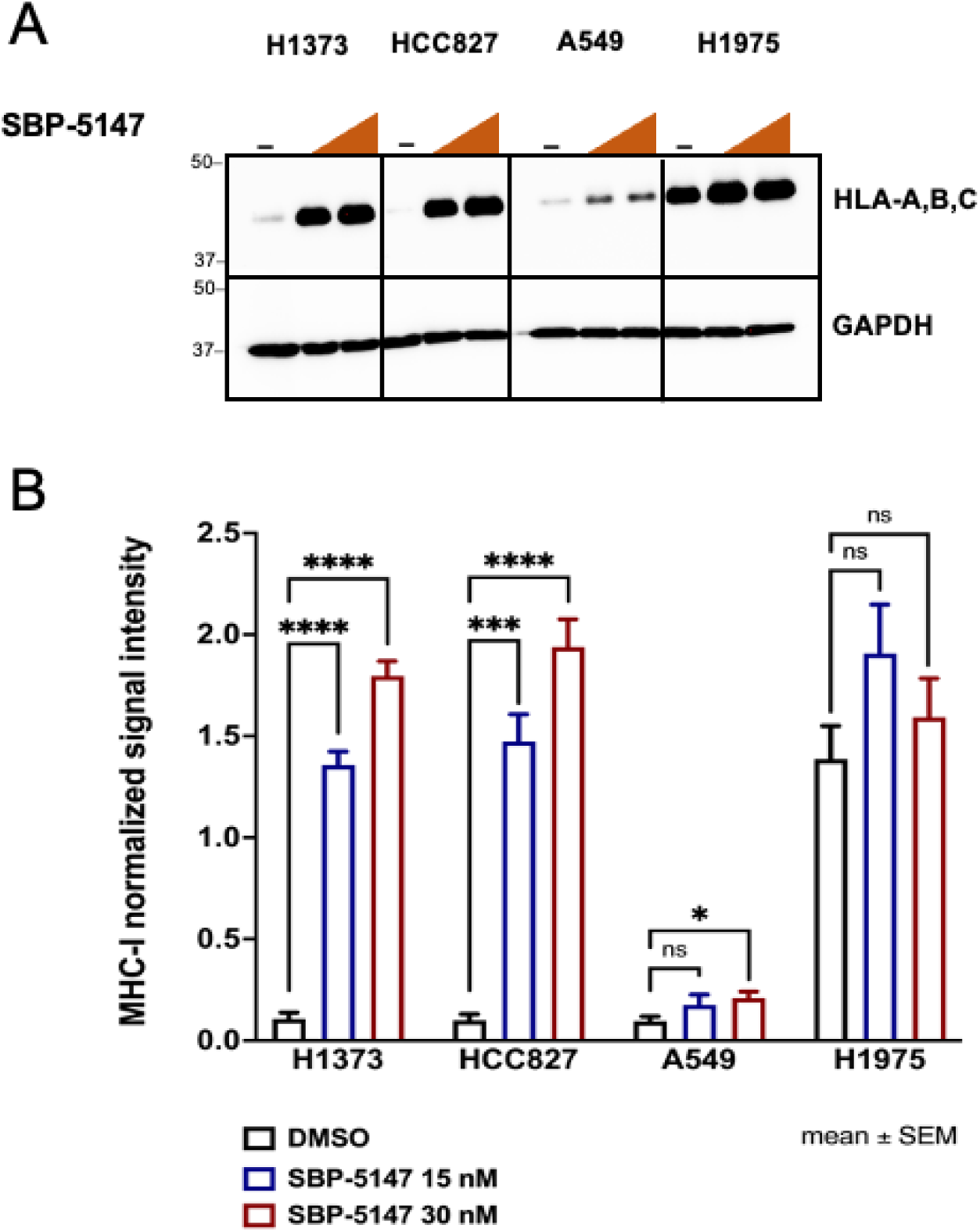
SBP-5147 increases levels of total MHC-I (HLA-A, -B, -C) protein. (**A**) NSCLC cells were treated with either DMSO or **SBP-5147** (15 & 30 nM) and incubated for 72 h at standard cell culture conditions. Subsequently, cellular lysates were immunoblotted with the indicated antibodies to evaluate the effect of **SBP-5147** treatment on total MHC-I protein in the indicated cell lines. A representative image of three independent experiments is shown. (**B**) The density of protein signals was quantified for DMSO and **SBP-5147**-treated samples and normalized to GAPDH. Data is shown as mean ± SEM. ****P<0.0001, ***P<0.001, *P<0.05, ns= P≥ 0.05 vs DMSO by one-tailed unpaired t test.

These data indicate that not only does **SBP-5147** kill NSCLC cells through archetypal apoptosis, but treatment with this inhibitor also upregulates MHC-I expression, which potentially sensitizes these cells to ICIs, thus promoting ULK1/2 inhibition as a therapeutic strategy to overcome tumor immune evasion and enhance responses to immunotherapy in lung cancer.

## CONCLUSIONS

Autophagy dysregulation in cancer plays a role in tumor progression and resistance to standard-of-care treatments. The discovery and development of small molecules that modulate different steps of the autophagy signaling cascade may increase the efficacy of current treatments ^9, 12^. As the only serine/threonine (S/T) kinase in the core autophagy pathway, ULK1 plays a central role in autophagy initiation during cellular stress ^27^ and has become an attractive target for drug development.

Our studies demonstrate that ULK1/2 inhibitor potency can be enhanced by designing analogs, such as compound **5**, that engage additional binding interactions with Asp102/95 of ULK1/2. We further improved cellular potency by modulating the lipophilicity of the inhibitors (see **Table S1**). Two analogs, **SBP-5147** and **SBP-7501,** displayed excellent cellular activity in the nanoBRET assay and showed improved PK and PD properties in mice compared with our earlier inhibitors. Both compounds effectively inhibited phosphorylation of ULK1/2 substrates Beclin-1 and Vps34 and significantly reduced autophagic flux in nutrient-challenged A549 cells. Pharmacokinetic analysis of **SBP-5147** in mice at 10 mpk demonstrated appreciable plasma levels maintained above the ULK1/2 NanoBRET IC_50_ for at least 8 h post-dosing. Moreover, **SBP-5147** induced a sustained decrease in ATG13 and ATG101 protein expression for up to 24 h in liver and lung, indicating robust on-target activity in tissues.

We are the first to show that ULK1/2 inhibitors reduce the expression of autophagy-related proteins for 24 h post-treatment in mouse tissues. Both **SBP-5147** and **SBP-7501** demonstrated efficacy at 24 h post-dose, an effect that persisted beyond plasma exposure, indicating a durable effect on this metabolic pathway in vivo. These findings suggest a novel mechanism for autophagy inhibition that involves not only kinase inhibition but also protein degradation. Consequently, these compounds may be suitable for once-a-day dosing in humans at clinically reasonable levels.

In NSCLC, disrupting autophagy initiation by selective ULK1 inhibition has been reported as a strategy to tackle autophagy-dependent cancer cell survival, tumor progression, and treatment resistance ^9, 21, 22, 27^. We demonstrated the cytotoxic potential of **SBP-5147** and **SBP-7501** as single agents against NSCLC cells, inducing cell death by archetypal apoptosis. Importantly, this cytotoxicity was not observed in non-tumorigenic lung cells, indicating a potential therapeutic window for cancer treatment. Our results support the rationale for ULK1/2 inhibition as an adjuvant strategy to improve responses to cancer treatment and overcome resistance, as studies have demonstrated that ULK1 inhibition re-sensitizes NSCLC cells to chemotherapeutics ^21^.

Moreover, given the interplay between autophagy signaling and immune regulation ^55^ ^8, 56^, we found that **SBP-5147** increases MHC class 1 expression in NSCLC cells. This result aligns with studies linking ULK1 inhibition to improved responses to immune checkpoint blockade in lung cancer ^22^. This potential role for ULK1/2 inhibition in enhancing antigen presentation is especially relevant in NSCLC, where broad downregulation of MHC-I ^55^ and impaired T cell-mediated anti-tumor responses ^1, 57^ are well documented. From a translational perspective, these novel ULK1/2 inhibitors could expand our understanding of autophagy dependence in cancer and the subtleties of autophagy modulation, particularly in combinatorial therapeutic strategies.

## EXPERIMENTAL SECTION

### General Chemistry Information

All reactions were performed in oven-dried glassware under an atmosphere of nitrogen with magnetic stirring. All solvents and chemicals used were purchased from commercial sources and used as received without further purification unless otherwise noted. Purity and characterization of compounds were established by a combination of LC-MS and NMR spectroscopy (established purity > 95% for all tested compounds). Proton (^1^H) carbon (^13^C) NMR and fluorine (^19^F) spectra were obtained on a JEOL400 spectrometer at 400 MHz, 101 MHz, and 376 MHz respectively. Chemical shifts are reported in δ (ppm) and internally referenced to residual deuterated solvent signals. The data for ^1^H NMRs are reported in terms of chemical shift (δ ppm), multiplicity, coupling constant (Hz), and proton integration. The data for ^13^C NMRs are reported in terms of chemical shift (δ ppm). HPLC-MS analyses were performed on a Shimadzu 2010EV LCMS using the following conditions: Kromasil C18 column (reverse phase, 4.6 mm x 50 mm); a linear gradient from 10% acetonitrile and 90% water to 95% acetonitrile and 5% water over 4.5 min; flow rate of 1 mL/min; and UV photodiode array detection from 200 to 300 nm. High-resolution ESI-TOF mass spectra were acquired from the Scripps Research Institute Center for Metabolomics and Mass Spectrometry in La Jolla, CA.

### General Method 1

#### 6-(4-Chloro-5-(trifluoromethyl)pyrimidin-2-ylamino)-3,4-dihydroquinolin-2(1H)-one (Synthesis of Int A, Step 1)

To a solution of 5-trifluromethyl-2,4-dichloropyrimidine (1 g, 4.6 mmol) in DCE:^t^BuOH (1:1, 40 mL) was added ZnCl_2_ (5.5 mL, 5.5 mmol) at 0 °C. After 1 h, 6-amino-3,4-dihydroquinolin-2(1H)-one hydrochloride (0.938 g, 4.6 mmol) and triethylamine (1 g, 5.5 mmol) in DCE:^t^BuOH (4 mL) was added to the reaction mixture. After stirring for 1.5 h, the reaction mixture was concentrated to get the crude product. The crude product was triturated with methanol and filtered and dried in vacuum to afford the compound (1 g, 64%) as a yellow solid. **^1^H NMR** (DMSO-*d_6_*) δ 10.51 (s, 1H), 10.04 (s, 1H), 8.79 (s, 1H), 7.40 (s, 1H), 7.39 (d, *J* = 8.2 Hz, 1H), 6.82 (d, *J* = 12.6 Hz, 1H), 2.76 (t, *J* = 7.3 Hz, 2H), 2.39 (t, *J* = 11.7 Hz, 2H). **^13^C NMR** (DMSO-*d_6_)*δ 170.6, 1601.2, 135.6, 133.4, 124.4, 121.2, 120.5, 115.6, 30.9, 25.6. LC-MS (ESI) calcd. For C_14_H_10_ClF_3_N_4_O [M+H]^+^: 343.04; found: 342.90.

#### Step 2

To a solution of 6-((4-chloro-5-(trifluoromethyl) pyrimidine-2-yl)amino)-3,4-dihydroquinolin-2(1*H*)-one (1.0 equiv.), appropriate amine or alcohol (1.0 equiv.) and *N, N*-diisopropylethylamine (3.0 equiv.) were mixed in acetonitrile or DMF and microwaved at 100 to 130 °C for 10 to 30 min. The reaction mixture was concentrated in vacuo and purified by trituration or automated chromatography to obtain the corresponding product.

### General Method 2

A solution of 2,4-dichloro-5-(trifluoromethyl)pyrimidine (1.0 equiv), the appropriate amine or alcohol (1.0 equiv), and *N, N*-diisopropylethylamine (1.0 equiv) in acetonitrile were microwaved at 100 °C for 10 min. The reaction mixture was concentrated in vacuo, and then the appropriate aniline (1.0 equiv) and acetic acid were added to the crude mixture and microwaved at 120 °C for 10 min. The reaction mixture was concentrated in vacuo and the crude product was purified by automated column chromatography to yield the corresponding product.

#### *N*^2^-Cyclopropyl-5-(trifluoromethyl)-*N*^2^-(3,4,5-trimethoxyphenyl)pyrimidine-2,4-diamine (1)

The title compound was prepared according to step 2, General Method 1 using commercially available 2-chloro-*N*-cyclopropyl-5-(trifluoromethyl) pyrimidine-4-amine (0.090 g, 0.379 mmol) and 3,4,5-trimethoxyaniline (0.069 g, 0.379 mmol). The crude product was purified by automated reverse phase chromatography. White solid (80 mg, 55%). LC-MS (ESI) calculated for C_17_H_20_F_3_N_4_O_3_ [M+H]^+^: 385.15; found 385.40. ^1^H NMR (DMSO-*d_6_*) δ 9.49 (s, 1H), 8.18 (s, 1H), 7.24 (s, 2H), 7.03 (d, *J* = 3.5 Hz, 1H), 3.72 (s, 6H), 3.62 (s, 3H), 3.05 – 2.96 (m, 1H), 0.76 – 0.65 (m, 4H). ^13^C NMR (DMSO-*d_6_*) δ 160.77, 159.34, 154.41 (q, *J* = 5.3 Hz), 152.64, 136.24, 132.85, 124.87 (q, *J* = 269.3 Hz), 97.66, 97.25 (q, *J* = 28.3 Hz), 60.15, 55.84, 24.47, 6.52. ^19^F NMR (DMSO-*d_6_*) δ −59.81. HRMS (ESI-TOF) calculated for C_17_H_20_F_3_N_4_O_3_ [M+H]^+^: 385.1482; found 385.1479.

#### 6-((4-(Cyclopropylamine)-5-(trifluoromethyl)pyrimidine-2-yl)amino)-3,4-dihydroquinoline-2(1H)-one (2; SBP-7501)

The title compound was prepared according to step 2, General Method 1 using 6-((4-chloro-5-(trifluoromethyl)pyrimidine-2-yl)amino)-3,4-dihydroquinoline-2(1*H*)-one (103 mg, 0.301 mmol), cyclopropylamine (21 µL, 0.301 mmol) and *N, N*-diisopropylethylamine (52 L, 0.301 mmol) were mixed in acetonitrile (3 mL) and microwaved at 120 °C for 10 min. It was concentrated in vacuo and purified by reverse phase HPLC to yield the product (90 mg, 82%). LC-MS (ESI) calculated for C_17_H_17_F_3_N_5_O [M+H]^+^: 364.14; found 364.36. ^1^H NMR (DMSO-*d_6_*) δ 9.96 (s, 1H), 9.56 (s, 1H), 8.15 (d, *J* = 0.9 Hz, 1H), 7.79 (s, 1H), 7.56 (d, *J* = 8.4 Hz, 1H), 7.10 (s, 1H), 6.75 (d, *J* = 8.6 Hz, 1H), 2.88 – 2.77 (m, 3H), 2.42 (dd, *J* = 8.3, 6.7 Hz, 2H), 0.76 (dt, *J* = 6.8, 3.2 Hz, 2H), 0.69 – 0.63 (m, 2H).^13^C NMR (DMSO-*d_6_*) δ 169.89, 160.72, 159.42, 154.47 (q, *J* = 5.5 Hz), 134.75, 132.80, 124.99 (q, *J* = 269.3 Hz), 123.50, 118.95, 118.21, 114.86, 30.51, 25.30, 24.48, 6.72, missing quartet Carbon at ±97 ppm. ^19^F NMR (DMSO-*d*_6_) δ −59.64. HRMS (ESI-TOF) calculated for C_17_H_17_F_3_N_5_O [M+H]^+^: 364.1385; found 364.1367.

#### 5-((4-(Cyclopropylamine)-5-(trifluoromethyl)pyrimidin-2-yl)amino)indolin-2-one (3)

2-Chloro-*N*-cyclopropyl-5-(trifluoromethyl) pyrimidine-4-amine (0.060 g, 0.253 mmol) and 5-aminoindolin-2-one (0.037 g, 0.253 mmol) were mixed in acetic acid (1 mL). The mixture was microwaved at 110 °C for 10 min. Filtered the solid and washed with acetonitrile to give 18 mg (20%) of the product. LC-MS (ESI) calculated for C_16_H_15_F_3_N_5_O [M+H]^+^: 350.12; found 350.10. ^1^H NMR (DMSO-*d*_6_) δ 10.51 (s, 1H), 10.39 (s, 1H), 8.33 (s, 1H), 8.12 (s, 1H), 7.71 (s, 1H), 7.50 (d, *J* = 7.2 Hz, 1H), 6.80 (d, *J* = 8.4 Hz, 1H), 3.49 (s, 2H), 2.87 (s, 1H), 0.84 – 0.76 (m, 2H), 0.73 (dq, *J* = 6.1, 4.1 Hz, 2H). ^13^C NMR (DMSO-*d*_6_) δ 176.28, 160.73, 159.35, 154.43 (q, J = 4.9 Hz), 138.22, 134.38, 125.77, 124.99 (q, *J* = 269.3 Hz), 118.55, 116.66, 108.64, 36.07, 24.49, 6.66. ^19^F NMR (DMSO-*d_6_*) δ −59.57. HRMS (ESI-TOF) calculated for C_16_H_15_F_3_N_5_O [M+H]^+^: 350.1229; found 350.1206.

#### *tert*-Butyl 6-((4-(cyclopropylamine)-5-(trifluoromethyl) pyrimidin-2-yl)amino)-3,4-dihydroisoquinoline-2(1*H*)-carboxylate (4)

2-Chloro-*N*-cyclopropyl-5-(trifluoromethyl) pyrimidine-4-amine (0.100 g, 0.421 mmol), diacetyl palladium (2.83 mg, 0.013 mmol), *tert*-butyl 6-amino-3,4-dihydroisoquinoline-2(1*H*)-carboxylate (0.115 g, 0.463 mmol) and cesium carbonate (0.178 g, 0.547 mmol) were mixed in 1,4-dioxane (2 mL) at room temperature and purged with nitrogen The mixture was microwaved at 130 °C for 20 min. The reaction mixture was filtered through a celite pad and the filtrate was concentrated under reduced pressure to afford the crude product which was purified by flash chromatography on silica gel (DCM-EtOAc) to afford the title compound. 166 mg (88%). LC-MS (ESI) calculated for C_22_H_26_F_3_N_5_O_2_+H]^+^:450.21, found; 450.55. ^1^H NMR (DMSO-*d*_6_) δ 9.65 (s, 1H), 8.18 (s, 1H), 7.84 (s, 1H), 7.61 (dd, *J* = 8.3, 2.2 Hz, 1H), 7.15 (s, 1H), 7.05 (d, *J* = 8.4 Hz, 1H), 4.42 (s, 2H), 3.53 (t, *J* = 5.9 Hz, 2H), 2.85 (dq, *J* = 7.3, 3.9 Hz, 1H), 2.72 (q, *J* = 6.4, 6.0 Hz, 2H), 1.41 (s, 9H), 0.79 (dt, *J* = 6.8, 3.3 Hz, 2H), 0.68 (p, *J* = 4.6 Hz, 2H). ^13^C NMR (DMSO-*d*_6_) δ 160.78, 159.44, 154.47 (q, *J* = 5.2 Hz), 154.05, 138.50, 134.44, 126.61, 126.20, 124.93 (q, *J* = 269.0 Hz), 118.85, 117.52, 97.12 (q, *J* = 32.2 Hz), 78.89, 45.19, 44.53, 41.62, 28.66, 28.12, 24.55, 6.72. ^19^F NMR (DMSO-*d*_6_) δ −59.81.

#### 4-Cyclopropyl-*N-2*(1,2,3,4-tetrahydroisoquinolin-6-yI)-5-(trifluoromethyl)pyrimidine-2,4-diamine\ HCl (5; SBP-5147)

*tert*-Butyl 6-((4-(cyclopropylamine)-5-(trifluoromethyl) pyrimidine-2-yl)amino)-3,4-dihydroisoquinoline-2(1*H*)-carboxylate (0.132 g, 0.294 mmol) and 3M hydrogen chloride in water (0.489 mL, 1.468 mmol) were mixed in methanol (1 mL). Heated at 60 °C for 16 h and then concentrated. The solid was washed with DCM to give 109 mg (96%) of the product. LC-MS (ESI) calculated for C_17_H_18_F_3_N_5_ [M+H]^+^: 350.16, found 350.05. ^1^H NMR (DMSO-*d*_6_) δ 9.78 (s, 1H, 8), 9.49 (s, 1H, 16), 8.17 (s, 1H, 3), 7.85 (s, 1H, 14), 7.60 (dd, *J* = 8.4, 1.8 Hz, 1H, 10), 7.25 (s, 1H, 7), 7.08 (d, *J* = 8.5 Hz, 1H, 11), 4.12 (t, *J* = 4.8 Hz, 2H, 15), 3.32 – 3.25 (m, 1H, 17), 2.92 (t, *J* = 6.0 Hz, 2H, 18), 2.85 – 2.78 (m, 1H, 20), 0.79 – 0.68 (m, 2H, 19’’, 25’’), 0.66 (q, *J* = 3.3, 2.4 Hz, 2H, 19’, 25’). ^13^C NMR (DMSO-*d*_6_) δ 160.75159.9, 154.49 (d, *J* = 5.4 Hz), 139.80,132.53, 127.37, 125.22 (q, *J* = 269.5 Hz), 122.66, 119.23, 118.43, 97.94 (q, J = 32.2 Hz), 43.83, 41.09, 25.56, 25.11, 7.21.^19^F NMR (DMSO-*d*_6_) δ −59.89. HRMS (ESI-TOF) calculated for C_17_H_18_F_3_N_5_ [M+H]^+^: 350.1587; found 350.1597.

#### 5-Bromo-N4-cyclopropyl-N2-(3,4,5-trimethoxyphenyl)pyrimidine-2,4-diamine (6)

5-Bromo-2,4-dichloropyrimidine (0.125 g, 0.549 mmol) and cyclopropanamine (0.038 mL, 0.549 mmol) were mixed in acetonitrile (2 mL) at 5 °C. After 2 min, *N, N*-diisopropylethylamine (0.096 mL, 0.549 mmol) was added and warmed to 21 °C. The mixture was microwaved at 70 °C for 10 min and then concentrated in vacuo. 3,4,5-trimethoxyaniline (0.100 g, 0.549 mmol) and acetic acid (2 mL) were added, and the reaction mixture was processed according to General Method 2. White solid (120 mg, 55%). LC-MS (ESI) calculated for C_16_H_20_BrN_4_O_3_ [M+H]^+^: 395.07; found 394.90. ^1^H NMR (DMSO-*d_6_*) δ 10.69 (s, 1H), 8.41 (s, 1H), 8.27 (s, 1H), 6.98 (s, 2H), 3.74 (s, 6H), 3.64 (s, 3H), 3.11 – 3.02 (m, 1H), 0.85 – 0.72 (m, 4H). ^13^C NMR (DMSO-*d_6_*) δ 160.28, 152.95, 151.47, 144.18, 134.47, 133.16, 99.10, 91.67, 60.18, 55.95, 25.42, 6.21. HRMS (ESI-TOF) calculated for C_16_H_20_BrN_4_O_3_ [M+H]^+^: 395.0719; found 395.0767.

#### 6-((5-Bromo-4-(cyclopropylamine)pyrimidin-2-yl)amino)-3,4-dihydroquinolin-2(1*H*)-one (7)

The title compound was prepared according to General Method 2 using 5-bromo-2,4-dichloropyrimidine (80 mg, 0.351mmol), cyclopropanamine (26 mg, 0.46 mmol) and *N, N*-diisopropylethylamine (0.092 mL, 0.53 mmol) in the first step, followed by 6-amino-3,4-dihydroquinolin-2(1*H*)-one (57 mg, 0.35 mmol) and acetic acid (2 mL) in the second step. 10 mg (7.6%) of the product was isolated by filtration of the crude solid and rinsing with acetonitrile. LC-MS (ESI) calculated for C_16_H_17_BrN_5_O [M+H]^+^: 374.06; found 374.26. ^1^H NMR (DMSO-*d*_6_) δ 9.87 (s, 1H, 17), 9.10 (s, 1H, 10), 7.93 (s, 1H, 3), 7.72 (s, 1H, 16), 7.46 (dd, *J* = 8.4, 1.8 Hz, 1H, 12), 6.95 (d, *J* = 3.3 Hz, 1H, 9), 6.68 (d, *J* = 8.7 Hz, 1H, 13), 2.81 – 2.71 (m, 3H, 8, 20), 2.37 (td, *J* = 7.4, 6.7, 2.1 Hz, 2H, 19), 0.76 – 0.67 (m, 2H), 0.64 – 0.57 (m, 2H). ^13^C NMR (DMSO-*d*_6_) δ 170.35 159.82 158., 156.45 136.04 132.58 123.96 118.79 118.03, 115.37, 92.20, 31.08,25.86, 24.86, 7.08. HRMS (ESI-TOF) calculated for C_16_H_17_BrN_5_O [M+H]^+^: 374.0616; found 374.0611.

#### 6-((4-(Cyclopropyl(methyl)amino)-5-(trifluoromethyl)pyrimidine-2-yl)amino)-3,4-dihydroquinolin-2(1*H*)-one (9)

The title compound was prepared according to step 2, General Method 1 using 6-((4-chloro-5-(trifluoromethyl) pyrimidine-2-yl)amino)-3,4-dihydroquinolin-2(1*H*)-one (75 mg, 0.219 mmol), *N*-methylcyclopropanamine-HC1 (24 mg, 0.219 mmol) and *N, N*-diisopropylethylamine (76 mL, 0.438 mmol) in acetonitrile (1 mL). The mixture was microwaved at 90 °C for 10 min and then concentrated in vacuo. The crude product was triturated with ethyl acetate and filtered to give the final product. Tan solid (40 mg, 48%). LC-MS (ESI) calculated for C_18_H_19_F_3_N_5_O [M+H]^+^: 378.15; found 378.30. ^1^H NMR (400 MHz, DMSO-*d_6_*): δ ^1^H NMR (DMSO-*d_6_*) δ 9.97 (s, 1H), 9.60 (s, 1H), 8.34 (s, 1H), 7.63 (s, 1H), 7.48 (dd, *J* = 8.6, 2.4 Hz, 1H), 6.75 (d, *J* = 8.6 Hz, 1H), 3.08 (s, 3H), 2.95 (tt, *J* = 7.3, 4.0 Hz, 1H), 2.81 (t, *J* = 7.5 Hz, 3H), 2.42 (dd, *J* = 8.5, 6.5 Hz, 2H), 0.79 (dd, *J* = 7.2, 5.3 Hz, 2H), 0.60 (p, *J* = 4.7 Hz, 2H). ^13^C NMR (DMSO-*d_6_*) δ 169.86, 161.89, 159.32, 157.78 (q, *J* = 5.1 Hz), 134.40, 132.97, 125.12 (q, *J* = 269.5 Hz), 123.50, 119.11, 118.34, 114.85, 99.12 (q, *J* = 32.3 Hz), 39.23, 33.97, 30.48, 25.23, 8.89. ^19^F NMR (DMSO-*d_6_*) δ −52.40. HRMS (ESI-TOF) calculated for C_18_H_19_F_3_N_5_O [M+H]^+^: 378.1542; found 378.1537.

#### 6-((4-Amino-5-(trifluoromethyl)pyrimidine-2-yl)amino)-3,4-dihydroquinoline-2(1*H*)-one (10)

A solution of 6-((4-chloro-5-(trifluoromethyl)pyrimidine-2-yl)amino)-3,4-dihydroquinoline-2(1*H*)-one (100 mg, 292 µmol), ammonia, 7 M THF (83.4 µL, 584 µmol) and DIPEA (56.6 mg, 438 µmol) in 5 mL DMF was heated at 100 °C for 2 h. The mixture was concentrated, triturated with methanol, and filtered. White solid (75 mg, 80%). LC-MS (ESI) calculated for C_14_H_13_F_3_N_5_O [M+H]^+^: 324.11; found 323.80. ^1^H NMR (DMSO-*d_6_*) δ 9.94 (s, 1H), 9.35 (s, 1H), 8.17 (s, 1H), 7.61 (d, *J* = 2.4 Hz, 1H), 7.52 (dd, *J* = 8.6, 2.4 Hz, 1H), 7.03 (bs, 2H), 6.76 (d, *J* = 8.6 Hz, 1H), 2.87 – 2.81 (m, 2H), 2.46 – 2.38 (m, 2H).^13^C NMR (DMSO-*d_6_*) δ 169.91, 161.13, 159.64, 155.07 (q, *J* = 4.5, 3.7 Hz), 134.63, 132.88, 125.16 (q, *J* = 269.0 Hz), 123.59, 119.38, 118.59, 114.89, 96.15 (q, *J* = 33.7 Hz), 30.53, 25.16. ^19^F NMR (DMSO-*d_6_*) δ −60.09. HRMS (ESI-TOF) calculated for C_14_H_13_F_3_N_5_O [M+H]^+^: 324.1072; found 324.1067.

#### 6-((4-(Cyclobutylamine)-5-(trifluoromethyl)pyrimidin-2-yl)amino)-3,4-dihydroquinolin-2(1*H*)-one (11)

The title compound was prepared according to General Method 2 using 2,4-dichloro-5-(trifluoromethyl)pyrimidine (70 mg, 0.32 mmol), cyclobutylamine (23 mg, 0.32 mmol) and *N, N*-diisopropylethylamine (0.056 mL, 0.32 mmol) in the first step, followed by 6-amino-3,4-dihydroquinolin-2(1*H*)-one (52 mg, 0.32 mmol) and acetic acid (2 mL) in the second step. The crude product was purified by automated reverse phase chromatography. White solid (12 mg, 15%). LC-MS (ESI) calculated for C_18_H_19_F_3_N_5_O [M+H]^+^: 378.15; found 378.35. ^1^H NMR (DMSO-*d_6_*) δ 9.97 (s, 1H), 9.48 (s, 1H), 8.15 (s, 1H), 7.67 (s, 1H), 7.38 (dd, *J* = 8.5, 1.9 Hz, 1H), 6.96 (d, *J* = 6.6 Hz, 1H), 6.77 (d, *J* = 8.5 Hz, 1H), 4.62 (q, *J* = 8.1 Hz, 1H), 2.85 (t, *J* = 7.6 Hz, 2H), 2.44 (dd, *J* = 8.4, 6.5 Hz, 2H), 2.31 – 2.20 (m, 2H), 2.19 – 2.10 (m, 2H), 1.75 – 1.59 (m, 2H). ^13^C NMR (DMSO-*d_6_*) δ 169.85, 160.77, 157.11, 154.77 (q, *J* = 5.5 Hz), 134.48, 132.99, 125.00 (q, *J* = 269.0 Hz), 123.45, 119.39, 118.56, 114.84, 96.60 (q, *J* = 38.2 Hz), 45.98, 30.50, 29.93, 25.27, 14.84. ^19^F NMR: δ −59.57. HRMS (ESI-TOF) C_18_H_19_F_3_N_5_O [M+H]^+^: 378.1536; found 378.1541.

#### 6-((4-(Cyclopentylamine)-5-(trifluoromethyl)pyrimidin-2-yl)amino)-3,4-dihydroquinolin-2(1*H*)-one (12)

The title compound was prepared according to General Method 2 using 2,4-dichloro-5-(trifluoromethyl)pyrimidine (120 mg, 0.55mmol), cyclopentanamine (47 mg, 0.55 mmol) and *N, N*-diisopropylethylamine (0.096 mL, 0.55 mmol) in the first step, followed by 6-amino-3,4-dihydroquinolin-2(1*H*)-one (90 mg, 0.55) and acetic acid (2 mL) in the second step. The crude product was purified by reverse phase HPLC. White solid (65 mg, 60%). LC-MS (ESI) calculated for C_19_H_21_F_3_N_5_O [M+H]^+^: 392.17; found 392.40. ^1^H NMR (DMSO-d6) δ 9.96 (s, 1H), 9.48 (s, 1H), 8.14 (s, 1H), 7.64 (s, 1H), 7.39 (dd, *J* = 8.7, 2.3 Hz, 1H), 6.75 (d, *J* = 8.6 Hz, 1H), 6.47 (d, *J* = 7.1 Hz, 1H), 4.53 – 4.41 (m, 1H), 2.82 (t, *J* = 7.6 Hz, 2H), 2.43 (t, *J* = 7.5 Hz, 2H), 2.01 – 1.88 (m, 2H), 1.76 – 1.65 (m, 2H), 1.65 – 1.47 (m, 4H). ^13^C NMR (DMSO-*d*_6_) δ 169.82, 160.78, 157.77, 154.66, 154.61, 154.56, 134.50, 132.97, 125.09 (q, *J* = 269.3 Hz), 123.42, 119.33, 118.54, 114.82, 96.68 (q, *J* = 35.8 Hz), 52.42, 40.15, 39.94, 39.73, 39.52, 39.31, 39.10, 38.89, 31.68, 30.49, 25.28, 23.51.^19^F NMR (DMSO-*d_6_*) δ −59.54. HRMS (ESI-TOF) calculated for C_19_H_21_F_3_N_5_O [M+H]^+^: 392.1698; found 392.1694.

#### 6-((4-(2-Ethyl Hydrazine Yl)-5-(trifluoromethyl)pyrimidin-2-yl)amino)-3,4-dihydroquinolin-2(1*H*)-one (13)

The title compound was prepared according to step 2, General Method 1 using 6-((4-chloro-5-(trifluoromethyl) pyrimidine-2-yl)amino)-3,4-dihydroquinolin-2(1*H*)-one (70 mg, 0.204 mmol), methylhydrazine-oxalic acid (31 mg, 0.204 mmol) and *N, N*-diisopropylethylamine (0.12 mL, 0.613 mmol) in acetonitrile (1 mL). The reaction mixture was microwaved at 100 °C for 10 min and then concentrated in vacuo. The product was triturated with acetone and filtered to afford the desired product. Tan solid (61 mg, 82%). LC-MS (ESI) calculated for C_16_H_18_F_3_N_6_O [M+H]^+^: 367.15; found 367.30. ^1^H NMR (DMSO-*d_6_*) δ 9.96 (s, 1H), 9.37 (s, 1H), 8.23 (s, 1H), 7.58 (d, *J* = 1.9 Hz, 1H), 7.36 (dd, *J* = 8.5, 2.3 Hz, 1H), 6.75 (d, *J* =8.5 Hz, 1H), 4.77 (s, 2H), 3.77 (q, *J* = 7.0 Hz, 2H), 2.82 (dd, *J* = 8.4, 6.5 Hz, 2H), 2.43 (dd, *J* = 8.6, 6.5 Hz, 2H), 1.21 (t, *J* = 7.1 Hz, 3H). ^13^C NMR (DMSO-*d_6_*) δ 169.85, 160.27, 159.47, 157.25 (q, *J* = 6.7 Hz), 134.67, 132.80, 125.40 (q, *J* = 266.8 Hz), 123.50, 119.03, 118.23, 114.86, 97.76 (q, *J* = 31.8 Hz), 47.65, 30.50, 25.24, 1057. ^19^F NMR (DMSO-*d_6_*) δ −51.34. HRMS (ESI-TOF) calculated for C_16_H_18_F_3_N_6_O [M+H]^+^: 367.1494; found 367.1486.

#### 6-((4-((2,2,2-Trifluoroethyl)amino)-5-(trifluoromethyl)pyrimidin-2-yl)amino)-3,4 dihydroquinolin-2(1*H*)-one (14)

The title compound was prepared according to Step 2, General Method 1 using 6-((4-chloro-5fz-(trifluoromethyl) pyrimidine-2-yl)amino)-3,4-dihydroquinolin-2(1*H)*-one (70 mg, 0.204 mmol), 2,2,2-trifluoroethanol-1-amine·HCl (28 mg, 0.204 mmol) and *N, N*-diisopropylethylamine (71 mL, 0.409 mmol) in DMF (1 mL). The reaction mixture was microwaved at 130 °C for 20 min and then concentrated in vacuo. The crude product was purified by automated reverse phase chromatography. White solid (22 mg, 27%). LC-MS (ESI) calculated for C_16_H_14_F_6_N_5_O [M+H]^+^: 406.11; found 406.30. ^1^H NMR (DMSO-*d_6_*) δ 9.99 (s, 1H), 9.67 (s, 1H), 8.27 (d, *J* = 0.9 Hz, 1H), 7.53 (d, *J* = 2.4 Hz, 1H), 7.34 (dd, *J* = 8.5, 2.4 Hz, 1H), 6.77 (d, *J* = 8.5 Hz, 1H), 4.24 (qd, *J* = 9.3, 6.1 Hz, 2H), 2.82 (dd, *J* = 8.5, 6.5 Hz, 2H), 2.46 – 2.39 (m, 2H). ^13^C NMR (, DMSO-*d_6_*) δ 169.88, 160.65, 158.29, 155.50 (q, *J* = 4.8 Hz), 133.93, 133.39, 124.94 (q, *J* = 280.6 Hz), 124.80 (q, *J* = 269.3 Hz), 123.52, 119.74, 119.00, 114.83, 97.15 (q, *J* = 26.3 Hz), 41.06 (q, *J* = 31.3 Hz), 30.43, 25.10. ^19^F NMR (DMSO-*d_6_*) δ −59.68, −69.92. HRMS (ESI-TOF) calculated for C_16_H_14_F_6_N_5_O [M+H]^+^: 406.1103; found 406.1093.

#### 6-((4-((2-Hydroxyethyl)amino)-5-(trifluoromethyl)pyrimidin-2-yl)amino)-3,4-dihydroquinolin-2(1*H*)-one (15)

The title compound was prepared according to step 2, General Method 1 using 6-((4-chloro-5-(trifluoromethyl) pyrimidine-2-yl)amino)-3,4-dihydroquinolin-2(1*H*)-one (70 mg, 0.204 mmol), 2-aminoethan-l-ol (12 mg, 0.204 mmol) and *N, N-* diisopropylethylamine (36 mL, 0.204 mmol) in acetonitrile (1 mL). The mixture was microwaved at 100 °C for 10 min and then concentrated in vacuo. The crude product was triturated with acetone and filtered to give the final product. Tan solid (53 mg, 71%). LC-MS (ESI) calculated for C_16_H_17_F_3_N_5_O_2_ [M+H]^+^: 368.13; found 368.10. ^1^H NMR (DMSO-*d_6_*) δ 9.97 (s, 1H), 9.48 (bs, 1H), 8.15 (s, 1H), 7.61 (s, 1H), 7.42 (dd, *J* = 8.6, 2.3 Hz, 1H), 6.86 (bs, 1H), 6.76 (d, *J* = 8.6 Hz, 1H), 4.78 (bs, 1H), 3.58 (dd, *J* = 7.7, 5.6 Hz, 2H), 3.52 (dd, *J* = 7.5, 6.4 Hz, 2H), 2.84 (dd, *J* = 8.5, 6.5 Hz, 2H), 2.42 (dd, *J* = 8.6, 6.5 Hz, 2H). ^13^C NMR (DMSO-*d_6_*) δ 169.92, 160.84, 158.22, 154.60 (q, *J* = 5.1 Hz), 134.50, 132.95, 125.16 (q, *J* = 269.1 Hz), 123.59, 119.28, 118.46, 114.92, 96.75 (q, *J* = 37.2 Hz), 59.10, 43.02, 30.52, 25.20. ^19^F NMR (DMSO-*d_6_*) δ −59.82. HRMS (ESI-TOF) calculated for C_16_H_17_F_3_N_5_O_2_ [M+H]^+^: 368.1334; found 368.1323.

#### 6-((4-((2-Methoxyethyl)amino)-5-(trifluoromethyl)pyrimidin-2-yl)amino)-3,4-dihydroquinolin-2(1*H*)-one (18)

The title compound was prepared according to General Method 2 using 2,4-dichloro-5-(trifluoromethyl)pyrimidine (70 mg, 0.323 mmol), 2-methoxyethan-l-amine (24 mg, 0.28 µL, 0.323 mmol) and *N, N-*diisopropylethylamine (56 mL, 0.323 mmol) in acetonitrile (2 mL) for the first step, followed by 6-amino-3,4-dihydroquinolin-2(1*H*)-one (52 mg, 0.323 mmol) in acetic acid (2 mL) for the second step. The crude product was purified by automated reverse phase chromatography. White solid (11 mg, 8.93%). LC-MS (ESI) calculated for C_17_H_19_F_3_N_5_O_2_ [M+H]^+^: 382.15; found 382.33.^1^H NMR (DMSO-*d*_6_) δ 9.97 (s, 1H), 9.49 (s, 1H), 8.15 (s, 1H), 7.59 (s, 1H), 7.42 (dd, *J* = 8.6, 2.4 Hz, 1H), 7.03 – 6.94 (m, 1H), 6.76 (d, *J* = 8.5 Hz, 1H), 3.60 (q, *J* = 6.1, 5.7 Hz, 2H), 3.50 (t, *J* = 6.2 Hz, 2H), 3.26 (s, 3H), 2.83 (dd, *J* = 8.5, 6.6 Hz, 2H), 2.43 (dd, *J* = 8.5, 6.4 Hz, 2H). ^13^C NMR (DMSO-*d*_6_) δ 169.88, 160.84, 158.16, 154.64 (q, *J* = 5.3 Hz), 134.46, 133.02, 125.11 (q, *J* = 269.3 Hz), 123.56, 119.37, 118.55, 114.87, 96.71 (q, *J* = 29.1 Hz), 69.93, 58.07, 40.02, 30.51, 25.16. 19F NMR (DMSO-*d_6_*) δ −59.87. HRMS (ESI-TOF) calculated for C_17_H_19_F_3_N_5_O_2_ [M+H]^+^: 382.1491; found 382.1491.

#### 6-((4-((3-Aminopropyl)amino)-5-(trifluoromethyl)pyrimidin-2-yl)amino)-3,4 dihydroquinolin-2(1*H*)-one·HCl (19)

*tert*-Butyl (3-((2-((2-oxo-1,2,3,4-tetrahydroquinoline-6-yl)amino)-5-(trifluoromethyl) pyrimidine-4-yl)amino)propyl)carbamate (0.232 g, 0.483 mmol) and aqueous hydrogen chloride (0.64 mL, 1.931 mmol) were mixed in methanol (2 mL). The reaction mixture was stirred at 80 °C then concentrated in vacuo to give the desired product. Tan solid (195 mg, 97%). LC-MS (ESI) calculated for C_17_H_20_F_3_N_6_O [M+H]^+^: 381.17; found 381.10. ^1^H NMR (, DMSO-*d_6_*) δ 10.02 (s, 1H), 9.54 (s, 1H), 8.44 (s, 1H), 8.15 (s, 1H), 7.58 (s, 1H), 7.41 (dd, *J* = 8.5, 2.3 Hz, 1H), 7.35 (bs, 1H), 6.78 (d, *J* = 8.5 Hz, 1H), 3.50 (s, 2H), 2.83 (t, *J* = 7.4 Hz, 2H), 2.76 (s, 2H), 2.43 (t, *J* = 7.5 Hz, 2H), 1.87 (s, 2H). ^13^C NMR (DMSO-*d_6_*) δ 169.92, 166.17, 160.90, 157.99, 154.60 (d, *J* = 5.9 Hz), 134.47, 133.06, 125.12 (q, *J* = 269.3 Hz), 123.56, 119.45, 118.67, 114.96, 96.91 (q, *J* = 38.3 Hz), 37.52, 36.54, 30.53, 27.02, 25.27. ^19^F NMR (DMSO-*d*_6_) δ −60.67. HRMS (ESI-TOF) calculated for C_17_H_20_F_3_N_6_O [M+H]^+^: 381.1651; found 381.1608.

#### *tert*-Butyl (3-((2-((2-oxo-1,2,3,4-tetrahydroquinoline-6-yl)amino)-5-(trifluoromethyl) pyrimidin-4-yl)amino)propyl)carbamate (20)

The title compound was prepared according to step 2, General Method 1 using 6-((4-chloro-5-(trifluoromethyl)pyrimidin-2-yl)amino)-3,4-dihydroquinolin-2(1*H*)-one (0.200 g, 0.584 mmol), tert-butyl 5-aminopentanoate (0.101 g, 0.584 mmol) and *N, N-*diisopropylethylamine (0.102 mL, 0.584 mmol) in DMF (5 mL). The mixture was microwaved at 100 °C for 20 min and then concentrated in vacuo. The crude product was purified by automated normal phase chromatography (7% MeOH in DCM). White solid (242 mg, 78%). LC-MS (ESI) calculated for C_22_H_28_F_3_N_6_O_3_ [M+H]^+^: 481.22; found 481.55. ^1^H NMR (DMSO-*d_6_*) δ 9.97 (s, 1H), 9.46 (s, 1H), 8.14 (s, 1H), 7.60 (d, *J* = 2.4 Hz, 1H), 7.40 (dd, *J* = 8.6, 2.4 Hz, 1H), 7.08 (t, *J* = 5.4 Hz, 1H), 6.84 (t, *J* = 5.9 Hz, 1H), 6.76 (d, *J* = 8.6 Hz, 1H), 3.43 (q, *J* = 6.4 Hz, 2H), 2.97 (q, *J* = 6.6 Hz, 2H), 2.83 (dd, *J* = 8.5, 6.5 Hz, 2H), 2.43 (dd, *J* = 8.5, 6.5 Hz, 2H), 1.68 (p, *J* = 6.7 Hz, 2H), 1.36 (s, 9H). ^13^C NMR (DMSO-*d_6_*) δ 169.85, 160.87, 158.00,155.73, 154.50 (q, *J* = 5.6 Hz), 134.50, 132.96, 125.15 (q, *J* = 269.2 Hz), 123.50, 119.30, 118.49, 114.88, 96.71 (q, *J* = 29.7 Hz), 77.56, 37.97, 37.34, 30.51, 28.97, 28.23, 25.26. ^19^F NMR (DMSO-*d_6_*) δ −60.09. HRMS (ESI-TOF) calculated for C_22_H_28_F_3_N_6_O_3_ [M+H]^+^: 481.2175.

#### *N*-(3-((2-((2-Oxo-1,2,3,4-tetrahydroquinolin-6-yl)amino)-5-(trifluoromethyl)pyrimidin-4-yl)amino)propyl)cyclopropanecarboxamide (21)

6-((4-((3-Aminopropyl)amino)-5-(trifluoromethyl) pyrimidine-2-yl)amino)-3,4-dihydroquinolin-2(1*H*)-one·HCl (18, 0.024 g, 0.058 mmol), cyclopropanecarbonyl chloride (5.22 μL, 0.058 mmol) and triethylamine (20 mL, 0.144 mmol) were mixed in DMF (1 mL). MeOH was added and stirred for 1 h, then concentrated in vacuo. The crude product was purified by automated normal phase chromatography (4% MeOH in DCM). White solid (19 mg, 74%). LC-MS (ESI) calculated for C_21_H_24_F_3_N_6_O_2_ [M+H]^+^: 449.19; found 449.00. ^1^H NMR (DMSO-*d_6_*) δ 9.97 (s, 1H), 9.47 (s, 1H), 8.14 (s, 1H), 8.06 (t, *J* = 5.7 Hz, 1H), 7.59 (d, *J* = 2.3 Hz, 1H), 7.43 (dd, *J* = 8.7, 2.3 Hz, 1H), 7.11 (d, *J* = 6.1 Hz, 1H), 6.77 (d, *J* = 8.5 Hz, 1H), 3.44 (q, *J* = 6.4 Hz, 2H), 3.11 (q, *J* = 6.6 Hz, 2H), 2.83 (t, *J* = 7.5 Hz, 2H), 2.43 (dd, *J* = 8.4, 6.6 Hz, 2H), 1.70 (p, *J* = 6.8 Hz, 2H), 1.48 (tt, *J* = 8.1, 4.7 Hz, 1H), 0.69 – 0.56 (m, 4H). ^13^C NMR (DMSO-*d*_6_) δ 172.80, 169.93, 160.90, 158.04, 154.55 (q, *J* = 4.9 Hz), 134.54, 132.99, 125.17 (q, *J* = 269.2 Hz), 123.56, 119.33, 118.52, 114.94, 96.76 (q, *J* = 30.6 Hz), 38.09, 36.31, 30.53, 29.03, 25.29, 13.68, 6.16. ^19^F NMR (DMSO-*d*_6_) δ −60.05. HRMS (ESI-TOF) calculated for C_21_H_24_F_3_N_6_O_2_ [M+H]^+^: 449.1913; found 449.1893.

### Crystallization and Structure Determination

The ULK1/2 kinase domains were purified and crystallized as previously described ^11^. X-ray data sets were collected at the Diamond Light Source (beamline I04) and the Swiss Light Source (beamline X06DA), and were processed and scaled with XDS and aimless, respectively ^58, 59^. Molecular replacement was performed with Phaser ^60^ using PDB entries 4WNO (ULK1) or 6QAV (ULK2) as search models. The structures were rebuilt in Coot and refined using REFMAC ^61, 62^. Data collection and refinement statistics are summarized in **Table S4**.

### In Silico Docking

Fully continuous flexible small molecule ligand docking to the ULK1 and ULK2 proteins, represented by grid interaction potentials, was conducted using the ICM docking algorithm as implemented in the ICM-Pro program (v3.9, Molsoft, LLC) as previously described ^2^. The coordinates of our new three-dimensional structures of ULK1/2 were converted into an ICM object, charges were assigned, orientations of sidechain amides were corrected, hydrogen atoms were added, and their positions optimized by energy minimization using MMFF. The docking receptor site was defined around the **SBP-5147** ligand. An energy-minimized three-dimensional molecular model of the compound was generated, and docking was performed using the implemented routine in ICM.

### ULK1/2 ADP-Glo Kinase Assay

The kinase reaction was performed at room temperature in 5 µL total volume containing 25 µM of ATP (Sigma-Aldrich #A7699). The ADP-Glo™ Reagent (5 µL, Promega #V912C) was added after incubating kinase and substrate with ATP to stop the kinase reaction and remove the remaining ATP. After incubation in room temperature for 30 min, 10 µL of the Kinase Detection Reagent (Promega #V917A) was added to convert ADP into ATP, generating a luminescent signal that correlates with kinase activity. Luminescence was read on a Tecan Spark® Multimode Microplate Reader. The reaction for the ULK1 ADP-Glo assay used 2 µg/mL of recombinant human ULK1 protein (1-649, SignalChem #U01-11G) and 80 µg/mL of myelin basic protein (MBP, Sigma-Aldrich #M1891). The ULK2 assay employed 4 µg/mL of recombinant human ULK2 protein (1-478, SignalChem #U02-11G) and 80 µg/mL of MBP. To determine IC_50_ values, compounds were tested in a 16-dose-response mode assay with 3-fold serial dilutions starting at 30 µM and Staurosporine (Fisher Bioreagents #BP2541100), a non-selective protein kinase inhibitor, was used as positive control.

### ULK1/2 NanoBRET Intracellular Target Engagement Assay

Human embryonic kidney cells (HEK293T) were obtained from ATCC (#CRL-3216™) and maintained in DMEM 1X (Gibco) media supplemented with 10% heat-inactivated fetal bovine serum (FBS, Gibco) and 1% Antibiotic-Antimycotic solution (Gibco) at standard cell culture conditions. Cells were transfected with luciferase NanoLuc-ULK1 Fusion Vector (Promega #NV2211) or NanoLuc-ULK2 Fusion Vector (Promega #NV2221) using the jetPRIME transfection reagent (Polyplus #101000046). Meanwhile, the assay plate (non-binding surface 384-well plate, Corning #3574BC) was spotted with test compounds serially diluted 1:3 or vehicle control (DMSO) using an Echo 555 Liquid Handler. After 24 h, complete NanoBRET 20X (20 µM) Tracer K-5 reagent (Promega #N2482), prepared in accordance with the manufacturer’s directions, was added to the assay plate (2 µL/well) following the addition of 18 µL/well of Opti-MEM™ I Reduced Serum Medium (Gibco #31985-062). The assay plate was sealed with a removable seal, shaken on an orbital shaker for 2 min, and centrifuged at 200 X g for 1 min. In the meantime, cells were trypsinized, spun down, and resuspended in Opti-MEM. Cells were seeded into the assay plate (4 × 10^3^ cells/well in 20 µL) and the plate was centrifuged for 30 s at 42 X g. The assay plate was incubated for 2 h at 37°C, 5% CO_2,_ and then brought to room temperature for 15 min. A 3X Complete Substrate Inhibitor Solution was prepared according to the manufacturer’s recommended ratios (a mixture of Opti-MEM, NanoBRET™ Nano-Glo® Substrate [Promega #N157A], and Extracellular NanoLuc® Inhibitor [Promega #N235A]) and added to the assay plate (20 µL/well). The plate was incubated at room temperature for 2 to 3 min and the donor (450 nm) and acceptor (610 nm) emission wavelengths were measured on a Tecan Spark® Multimode Microplate Reader.

### Protein Thermal Shift Assay

Recombinant kinase domains of ULK2 at 2 µM in assay buffer (10 mM HEPES-NaOH, pH 7.5, 500 mM NaCl) were mixed with 1X SYPRO orange (Invitrogen #S6651) and 10 µM of ULK1/2 inhibitors. A temperature scan was run from 25°C to 95°C at a rate of 3°C/min. Temperature-dependent protein unfolding profiles (protein thermal shifts) were measured on a Real-Time PCR Mx3005p thermal cycler (Stratagene). The transition midpoint, in which the concentration of folded protein is equivalent to the concentration of unfolded protein, was defined as the melting temperature (*T_m_*). Data were analyzed using Protein Thermal Shift™ v1.4 (Thermo Fisher Scientific).

### phospho-Beclin-1 Assay

HEK293T cells were seeded into tissue culture-treated 6-well plates (Thermo Scientific #140675) at a density of 6 × 10^5^ cells/well (in 2.5 mL) and maintained in DMEM 1X (Gibco) media supplemented with 10% heat-inactivated FBS (Gibco) and 1% Antibiotic-Antimycotic solution (Gibco) at standard cell culture conditions. The day after cells were transfected with Myc-tagged kinase inactive (KI) ULK1 or wildtype (WT) ULK1 plus WT Flag-tagged Beclin-1 using the jetPRIME transfection reagent (Polyplus #101000046). At 24 h post-transfection, fresh media containing either DMSO or ULK1/2 inhibitors (10 µM) was added to the cells for 1 h. Next, cellular lysates were isolated in RIPA lysis buffer (Thermo Scientific #89900) containing protease and phosphatase inhibitors (Thermo Scientific #A32959). Lysates were immunoblotted with phospho-Beclin-1 Ser15 (Abbiotec #254515) and Myc-Tag (71D10) (Cell Signaling Technology [CST] #2278) antibodies, and membranes were visualized on a ChemiDoc Imaging System (Bio-Rad). Subsequently, membranes were stripped with Restore™ Western Blot Stripping Buffer (Thermo Scientific #21059) and incubated with Beclin-1 antibody (CST #3738) to probe total Beclin-1. GAPDH (14C10) antibody (CST #2118) was used as loading control. Bio-Rad Image Lab software 6.1 was used for densitometry analyses with background subtraction at 50.0 mm (disk size) and band detection sensitivity at 50%. Phospho-Beclin-1 Ser15, Beclin-1, and GAPDH protein signal densities (i.e., adjusted total band volume) were quantified for DMSO and compound-treated samples.

### phospho-Vps34 Assay

HEK293T cells were seeded and maintained as described for phospho-Beclin-1 assay. The next day cells were transfected with Myc-tagged kinase inactive (KI) ULK1 or WT ULK1 plus WT Flag-tagged Vps34 using Lipofectamine™ 2000 Transfection Reagent (Invitrogen #11668500), in conformity with its standard protocol. At 24 h post-transfection, fresh medium containing either DMSO or ULK1/2 inhibitors (10 µM) was added to the cells for 1 h. Next, cells were lysed with RIPA lysis buffer (Thermo Scientific #89900) containing protease and phosphatase inhibitors (Thermo Scientific #A32959). Cell lysates were immunoblotted with phospho-PI3 Kinase Class III (Ser249) (p-Vps34) (CST #13857), Myc-Tag (71D10) (CST #2278), and anti-FLAG® polyclonal (Sigma-Aldrich #F7425) antibodies. Membranes were visualized on a ChemiDoc Imaging System (Bio-Rad) and GAPDH (14C10) antibody (CST #2118) was used as loading control. Phospho-Vps34 (Ser249), Flag-Vps34, and GAPDH protein densities were quantified for DMSO and compound-treated samples, using Bio-Rad Image Lab software 6.1. Densitometry analyses were performed as described for the phospho-Beclin-1 assay.

### mCherry-GFP-LC3 FACS Assay

A549 cells (ATCC #CCL-185™) were transduced with the reporter pBABE-N-mCherry-GFP-LC3 and seeded into tissue culture-treated 6-well plates (Thermo Scientific #140675) at a density of 6 × 10^5^ cells/well (in 2.5 mL). Cells were maintained in enriched media (DMEM/F-12, GlutaMax ™ [Gibco #10565018] + 10% heat-inactivated FBS [Gibco]+ 1% Antibiotic-Antimycotic solution [Gibco]), starvation media (Sigma-Aldrich Earle’s balanced salt solution [EBSS] #E7510), or starvation media plus ULK1/2 inhibitors (10 µM) for 18 h. Subsequently, cells were trypsinized and resuspended in Phosphate Buffered Saline (PBS, Corning® #21-040). Analytical cytometry was performed at the Flow Cytometry Core of Sanford Burnham Prebys Medical Discovery Institute using an LSRFortessa 14-color (BD Biosciences) with a 488 nm and 610 nM laser. Wildtype A549 cells were used as unstained control, while A549 cells expressing either mCherry or GFP were used for compensation. 10,000 events were collected for each sample.

### Cell Viability Assay

The following non-small cell lung cancer cells (NSCLC) were obtained from ATCC: A549 (#CCL-185™), H1373 (#CRL-5866™), H1975 (#CRL-5908™), and HCC827 (#CRL-2868™). BEAS-2B, a non-tumorigenic epithelial cell line established from human bronchial epithelium, was also acquired from ATCC (#CRL-9609™). Cells were maintained in RPMI 1640 media with L-Glutamine (Corning) supplemented with 10% heat-inactivated FBS (Gibco) and 1% Antibiotic-Antimycotic solution (Gibco) at standard cell culture conditions. Cells were seeded (1,000 cells in 20 µL) into 384-well plates (#781098, Greiner) spotted with test compounds **SBP-5147** and **SBP-7501** (3-fold serial dilutions, from 30 µM to 1.5 nM), vehicle control (DMSO), or staurosporine (positive control) (Fisher Bioreagents #BP2541100) using an Echo 555 Liquid Handler. Cells were incubated for 72 h at standard cell culture conditions. Cell viability was assessed using the ATP-depletion assay CellTiter-Glo® (Promega #G7573) by adding 10 µL of its reagent mix to each well, as well as incubating the plate at room temperature for 10 min. Luminescence was read on a Tecan Spark® Multimode Microplate Reader (integration time= 1 s) and data was analyzed using GraphPad Prism software v10.3.0. The percentages compared to DMSO vehicle control were curve-fitted using nonlinear regression (log [inhibitor] vs. response, variable slope, four parameters).

### Pharmacokinetic (PK) and Pharmacodynamic (PD) Studies

All experimental protocols were approved by the Institutional Animal Care and Use Committee (IACUC) of Sanford Burnham Prebys Medical Discovery Institute and performed in conformity with regulations and guidelines of the National Institutes of Health (NIH) and the Association for Assessment and Accreditation of Laboratory Animal Care (AAALAC). Eight-week-old female C57BL/6J mice (The Jackson Laboratory #000664) were housed in groups of three in ventilated cages. The environmental conditions included temperature ∼25°C, 40%-70% humidity, and a 12-h light/12-h dark cycle. All animal procedures were performed during the light period of the cycle. The mice were fed a standard laboratory diet and food and water were supplied ad libitum. Mice were dosed with **SBP-5147**, **SBP-7501**, or vehicle (5% DMSO, 10% Tween 80, and 85% distilled and deionized H_2_O) via oral gavage (PO, 10 mg/kg). Both compounds were formulated in the described vehicle solution. At 15-30 min and 1, 2, 4, 8, and 24 h post dosing, blood samples were collected retro-orbitally, and plasma was separated by centrifugation. Plasma samples were extracted with acetonitrile: water (4:1) with 0.1% formic acid containing indomethacin as internal standard. Samples were centrifuged and supernatants were diluted with acetonitrile: water (4:1) with 0.1% formic acid. Compound concentration was determined by LC-MS/MS on a Shimadzu Nexera X2 HPLC coupled to an AB Sciex 6500 QTRAP, and data was analyzed using PKSolver software. For the PD studies, mice were dosed with **SBP-5147** (PO, 10 mg/kg) or vehicle as described above and sacrificed at 2-, 4-, 8-, and 24-h post-dosing. Liver and lung samples were flash frozen in liquid nitrogen and in vivo target engagement was assessed by immunoblotting. Total protein extracts were prepared in T-PER™ Tissue Protein Extraction Reagent (Thermo Scientific, #78510) containing a protease inhibitor cocktail (Thermo Scientific #A32953). The lysates were immunoblotted with Atg13 (D4P1K) (CST #13273) and Atg101 (E1Z4W) (CST #13492) antibodies. GAPDH (14C10) antibody (CST #2118) was used as loading control. Images were obtained on a ChemiDoc Imaging System (Bio-Rad) and Bio-Rad Image Lab software 6.1 was used for densitometry analyses with background subtraction at 50.0 mm (disk size) and band detection sensitivity at 50%. Atg13, Atg101, and GAPDH protein signal densities (i.e., adjusted total band volume) were quantified for vehicle control and experimental samples.

### Cell Death Assays

A nucleic acid stain strategy was used to measure the kinetics of cell death, in which cell viability is determined by means of exclusion of the DNA-binding fluorophore Sytox Green. One day prior to the experiment, A549 (ATCC #CCL-185™) and H1975 (ATCC #CRL-5908™) cells were seeded into 96-well plates (Perkin Elmer #6055302) at a density of 5,000 cells per well. The culture media was replaced with FluoroBrite DMEM (Gibco #A1896701) containing 10% heat-inactivated FBS and supplemented with 2 mM L-glutamine, 100 IU/mL penicillin, and 100 µg/mL streptomycin (Corning). Inhibitors targeting the major modalities of regulated cell death were utilized to determine which mechanisms are triggered by ULK1/2 inhibition. Cells were incubated for 1 h with 20 µM zVAD-FMK (Enzo #ALX-260-020), 20 µM Emricasan (IDN-6556, MedKoo #510230), 50 µM VX-765 (MedKoo #203165), 20 µM MCC950 (MedKoo #522637), 50 µM Necrostatin-1 (MedKoo #522437), or 20 µM Necrosulfonamide (MedKoo #525389). Upon incubation, cells were treated with 1 µM Hoechst staining (Thermo Scientific #33342) and 1.5 µM Sytox Green (Invitrogen #S7020). Subsequently, cells were treated with **SBP-5147** (1 µM), **SBP-7501** (5 µM), or DMSO and imaged for 24 h. Fluorescent cells were imaged and quantitated using a Cytation 5 plate reader (Biotek) at 10X magnification with the 377/447 nm and 469/525 nm filters. Four pictures per well were averaged every hour for 24 h. The total cell number was determined by Hoechst staining, while dead cells were identified by Sytox Green staining. In addition, cell death mechanisms were verified by immunoblotting of cell death-related proteins. Cells were seeded at 80% confluency in 3.5 cm diameter dishes a day prior to the experiment. Cells were pre-treated with cell death inhibitors and ULK1/2 inhibitors as described above. Cell lysates were isolated in RIPA buffer (RPI #R26200) containing protease inhibitors, a phosphatase inhibitor cocktail (CST #5870), and 2.5 µM biotin-ahx-DEVD-AOMK caspase activity-based probe. The following protease inhibitors were used: 20 µM 3,4-dichloroisocoumarin (DCI) (Calbiochem #287815), 10 µM E-64 (Sigma #E3132), 10 µM pepstatin A (Fisher #BP2671), and 10 µM MG-132 (Enzo #89161568). Cell lysates were immunoblotted with anti-PARP (CST #9532), anti-caspase-3 (Abcam #184787), and anti-gasdermin D (Novus #NBP2-33422) antibodies. Membranes were visualized on an ODYSSEY CLx imager (LI-COR). Pyroptosis was induced in THP-1 cells (ATCC #TIB-202™) via incubation with 10 µg/mL lipopolysaccharide (LPS, InvivoGen #tlrl-3pelps) for 4 h followed by 25 µM Nigericin (EMD #481990) for 2 h, with or without 20 µM MCC950. THP-1 cell lysates were utilized as positive control for gasdermin D cleavage, while β-tubulin (CST #86298) antibody was used as loading control.

### Total MHC-I Expression Assay

A549 (ATCC #CCL-185™), H1373 (ATCC #CRL-5866™), H1975 (ATCC #CRL-5908™), and HCC827 (ATCC #CRL-2868™) cells were maintained in RPMI 1640 media with L-Glutamine (Corning) supplemented with 10% heat-inactivated FBS (Gibco) and 1% Antibiotic-Antimycotic solution (Gibco) at standard cell culture conditions. Cells were seeded into tissue culture-treated 6-well plates (Thermo Scientific #140675) at a density of 1.5 × 10^5^ cells/well (in 3 mL) and incubated overnight at standard cell culture conditions. The next day cells were treated with either DMSO or **SBP-5147** (15 & 30 nM) for 72 h. Subsequently, cellular lysates were isolated in RIPA lysis buffer (Thermo Scientific #89900) containing protease and phosphatase inhibitors (Thermo Scientific #A32959). Lysates were immunoblotted with Anti-HLA Class 1 ABC (EMR8-5) (Abcam #ab70328) antibody and GAPDH (14C10) antibody (CST #2118) was used as loading control. Membranes were visualized on a ChemiDoc Imaging System (Bio-Rad) and Bio-Rad Image Lab software 6.1 was used for densitometry analyses with background subtraction at 50.0 mm (disk size) and band detection sensitivity at 50%. MHC-I and GAPDH protein signal densities (i.e., adjusted total band volume) were quantified for DMSO and compound-treated samples.

### Immunoblotting

Protein concentrations were determined by Pierce™ BCA Protein Assay Kit (Thermo Scientific #23227). Equal amounts of protein were separated on Novex™ Tris-Glycine 4-20% or 16% gels (Thermo Scientific #XP04200BOX, #XP04205BOX), or Bolt™ 4-12% Bis-Tris gels (Invitrogen #NW04122BOX) by SDS-PAGE and transferred to 0.2 µm nitrocellulose (Bio-Rad #1620168) or 0.2 µm PVDF membranes (Bio-Rad #1620174). Membranes were blocked in 5% dry milk (AmericanBio #AB10109-01000) or 5% BSA (Sigma-Aldrich #A7906-50G) in TBS-Tween 20 (0.1% vol/vol; TBS-T) for 1 h at room temperature. Protein-bound membranes were incubated with indicated primary antibodies in the same buffer overnight at 4°C. The next day membranes were washed three times with TBS-T (3 x 15 min) and then incubated for 1 h with horseradish peroxidase-conjugated secondary antibodies: anti-mouse IgG (CST# 7076S) or anti-rabbit IgG (CST #7074V). The following secondary antibodies were used for membranes imaged on an ODYSSEY CLx imager (LI-COR): IRDye 680 donkey anti-rabbit (LI-COR #926-68037), IRDye 800 donkey anti-mouse (LI-COR #926-32212), or IRDye® 800CW Streptavidin (LI-COR #926-32230) for probe detection. Membranes were washed 3 x 15 min with TBS-T prior to scanning.

### Statistical Analysis

Data analyses were performed using GraphPad Prism 10.3.0 software (GraphPad Software, La Jolla, CA, USA). Results are expressed as mean ± SEM or mean ± SD unless otherwise indicated. Student’s t-test or ANOVA were used to evaluate differences between experimental groups. The 50% inhibitory concentrations (IC_50_ values) were obtained by nonlinear regression analysis (log [inhibitor] vs. response, variable slope, four parameters).

## ASSOCIATED CONTENT

## Data Availability

The coordinates and structure factors of the ULK1 and ULK2 kinase domains in complex with compound SBP-5147 have been deposited in the Protein Data Bank (PDB) under accession numbers 9SE8 and 9SE9, respectively.

## Supporting Information

A Figure showing the structures of published ULK1/2 inhibitors, a schematic showing the principle of ADP-Glo, NanoBRET, and protein thermal shift assays, a Table showing the ratio of in vitro ULK1 activity (ADP-glo) versus in cell target engagement (NanoBRET) compared to ClogP values, a Table showing ULK1 and ULK2 ADP-Glo and NanoBRET data of selected compounds, a Table of ULK2 protein thermal shift data of selected compounds, a Figure showing the interaction of ULK1 inhibitors with the Asp102 side chain, a Figure showing ULK1/2 inhibitors induce apoptosis in H1975 cells, and a Table of the data collection and refinement statistics for ULK1 and ULK2 structures.

## AUTHOR INFORMATION

## Author Contributions

The manuscript was written through contributions of all authors. All authors have given approval to the final version of the manuscript.

## Notes

The authors declare no competing financial interest.

## ACKNOWLEDGMENTS

This work was supported by The Epstein Family Foundation, Pancreatic Action Network Foundation Grant 19 65 COSF, 2022 Curebound Discovery Grant, and 2023 CureBound Discovery Grant 23TG05. Research reported in this publication was supported by the National Cancer Institute of the National Institutes of Health under Award Number P30CA030199. The authors would like to thank the staff of the SBP Shared Resources Flow Cytometry Core and Animal Facility for their support in experimental design and analyses under this Cancer Center Support Grant. The content herein is solely the responsibility of the authors and does not necessarily represent the official views of the National Institutes of Health. SK and AC are grateful for support by the SGC, a registered charity that receives funds from Bayer AG, Boehringer Ingelheim, Bristol Myers Squibb, Genentech, Genome Canada, EU/EFPIA/OICR/McGill/KTH/Diamond Innovative Medicines Initiative 2 Joint Undertaking [EUbOPEN grant 875510], Janssen, Pfizer and Takeda. S.K. would like to acknowledge funding from the Frankfurt Cancer Institute (FCI), an institute supported by LOEWE and for support by the German Cancer Aid translational grant TACTIC.

## ABBREVIATIONS

ADME: absorption, distribution, metabolism, and excretion
ADP: Adenosine diphosphate
AMPK: adenosine monophosphate-activated protein kinase
APM: antigen presentation machinery
ATG: autophagy
ATP: Adenosine triphosphate
BRET: bioluminescence resonance energy transfer
DCM: dichloromethane
DIPEA: *N, N*-diisopropylethylamine
DMF: dimethylformamide
DMSO: dimethyl sulfoxide
EGFR: epidermal growth factor receptor
Eq: equivalent
FIP200: focal adhesion kinase family interacting protein of 200 kDa
FRET: fluorescent energy transfer
GAPDH: Glyceraldehyde 3-phosphate dehydrogenase
GFP: green fluorescent protein
HLA: Class 1 ABC, human leukocyte antigen class 1-A, -B, and -C
HRMS (ESI-TOF): high-resolution mass spectrometry (electrospray ionization process-time-of-flight)
HPLC: High-performance liquid chromatography
ICM: internal coordinate mechanics
IP: intraperitoneal
LC-MS (ESI): liquid chromatography-mass spectrometry (electrospray ionization process)
LC3: microtubule-associated protein 1 light chain 3 (MAP1LC3)
LKB1: liver kinase B1
MHC-I: major histocompatibility complex type 1
MMFF: Merck molecular force field
mTOR: mechanistic target of rapamycin
NMR: nuclear magnetic resonance
NSCLC: non-small cell lung cancer
PARP: poly ADP-ribose polymerase
PD: pharmacodynamic
PDB: protein data bank
PK: pharmacokinetics
PTS: protein thermal shift
QTRAP: quadrupole/linear ion trap mass spectrometer
SAR: structure-activity relationship
TBS-T: tris-buffered saline-tween 20
THF: tetrahydrofuran
TKI: tyrosine kinase inhibitor
ULK: *unc-51*-like autophagy activating kinase
VPS34: vacuolar protein sorting 34
WT: wild type

