## Supplemental Information for "Synthesis and Characterization of ULK1/2 Kinase Inhibitors that Inhibit Autophagy and Upregulate Expression of Major Histocompatibility Complex I for the Treatment of Non-Small Cell Lung Cancer"

^1^Center for Therapeutics Discovery, NCI-Designated Cancer Center, Sanford Burnham Prebys Medical Discovery Institute, La Jolla, California 92037, United States. ^2^Structural Genomics Consortium, Buchmann Institute for Molecular Life Sciences, Goethe-University Frankfurt, Frankfurt 60438, Germany. ^3^Institute of Pharmaceutical Chemistry, Goethe-University Frankfurt, Frankfurt 60438, Germany. ^4^Molecular and Cell Biology Laboratory, The Salk Institute for Biological Studies, La Jolla, California 92037, United States

*Co-first Authors **Corresponding Author

**SUPPORTING INFORMATION**

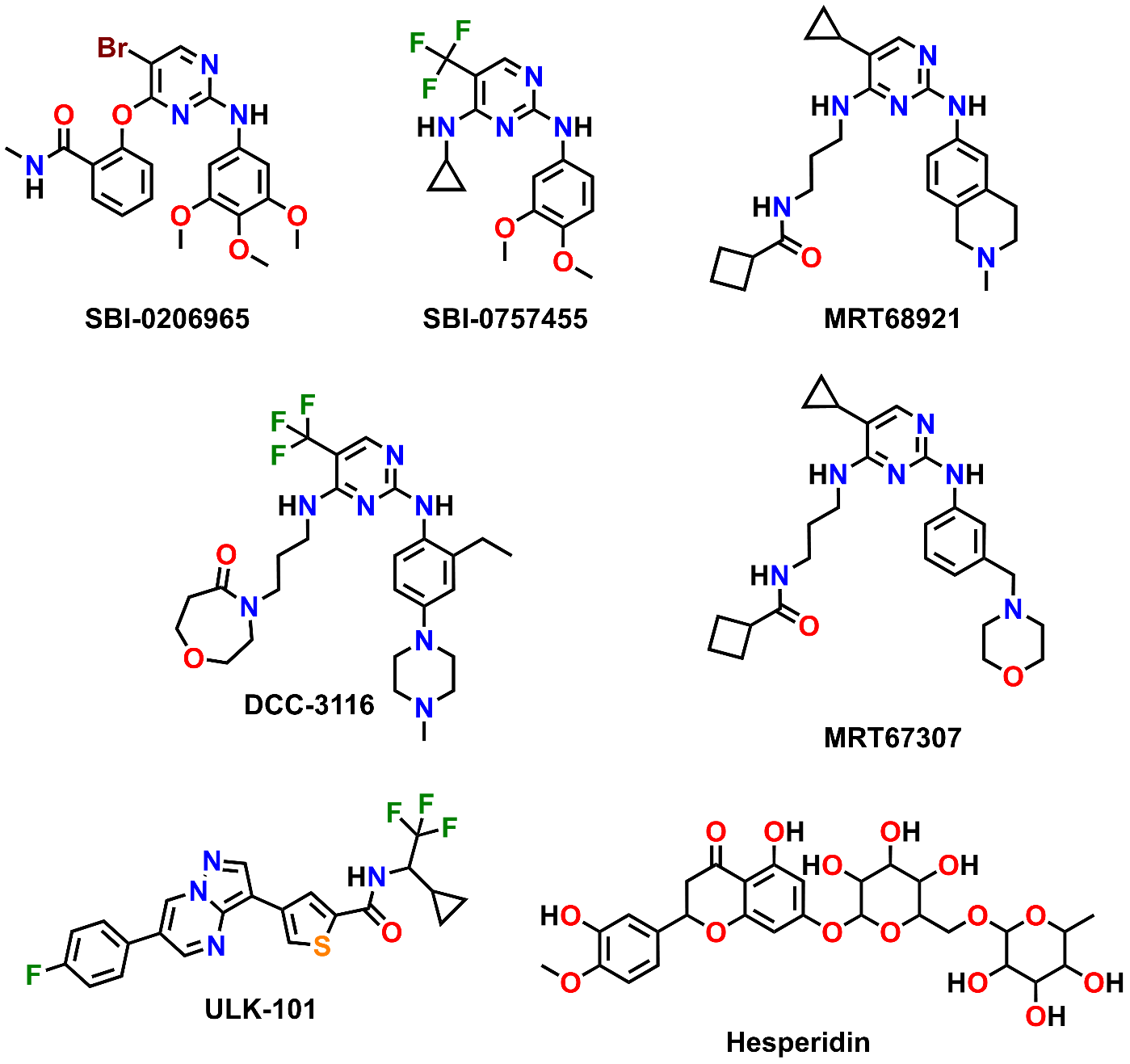

**Figure S1. Structures of ULK1/2 inhibitors.** **SBI-0206965** (Egan et al., 2015), a first-generation ULK1 inhibitor, **SBP-7455** (Ren et al., 2020), a dual ULK1/2 small-molecule inhibitor, both published by the Cosford laboratory at SBP. Other ULK inhibitors targeting autophagy in cancer include MRT68921 (Petherick et al., 2015), DCC-3116 (Ghazi et al., 2024), MRT67307 (Petherick et al., 2015), ULK-101 (Martin et al., 2018), and Hesperidin (Chaikuad et al., 2019).

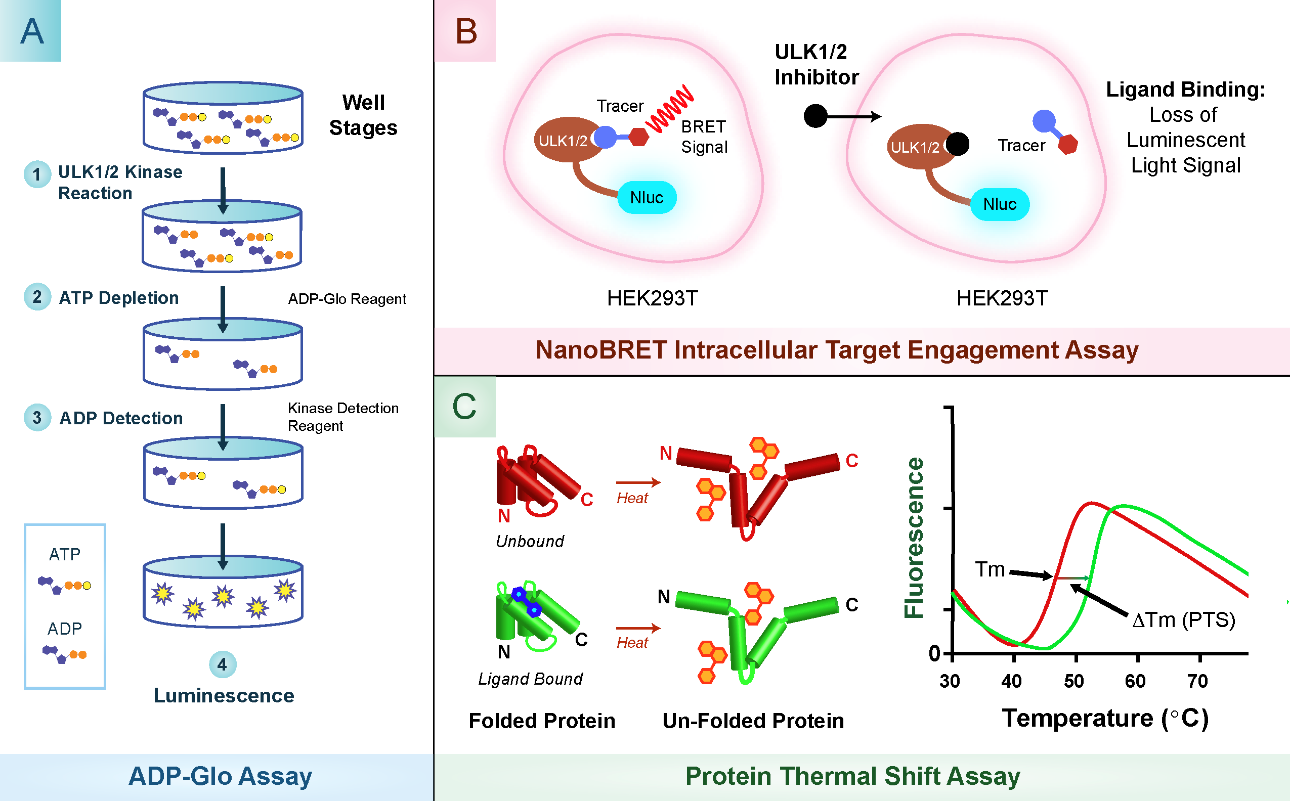

**Figure S2. Principle of ADP-Glo, NanoBRET, and Protein Thermal Shift Assays.** (**A**) Principle of the ADP-Glo Assay. The ADP-Glo Reagent is added after incubating kinase and substrate with ATP to stop the kinase reaction and deplete the remaining ATP. The Kinase Detection Reagent is then added to convert the ADP generated from the kinase reaction to ATP and allow the quantification of the ATP via luciferase/luciferin reaction. The luminescent signal is proportional to the amount of ADP generated during the kinase reaction, which is a measure of the kinase activity. Scheme from Promega. (**B**) Principle of the NanoBRET Intracellular Target Engagement Assay. ULK1 or ULK2 proteins are tagged with a luciferase, NanoLuc, and are transfected into HEK293T cells. A small molecule fluorescent tracer binds to the kinase and generates a BRET signal. The introduction of ULK1/2-binding inhibitors competitively displaces the tracer, resulting in a dose-dependent decrease of the NanoBRET signal, which allows quantification of intracellular binding affinity. Scheme from Promega. (**C**) Principle of the Protein Thermal Shift Assay. This assay is based on differential scanning fluorimetry in which a fluorescent dye has affinity for hydrophobic sites on proteins. As the temperature increases, the protein unfolds and the fluorescent dye binds to the exposed hydrophobic sites of the unfolded protein, leading to an increase in fluorescence. The transition midpoint, in which the concentration of folded protein equals the concentration of unfolded protein, is defined as the melting temperature (*Tm*). Ligands that bind and stabilize a protein will cause an increase of *Tm* that is proportional to the ligand’s affinity. Scheme adapted from Niesen et al. (Niesen et al., 2007).

**Table S1. Ratio of in vitro ULK1 activity (ADP-glo) versus in cell target engagement (NanoBRET) compared to ClogP values.**

| **Compound ID** | **Ratio of ULK1 ADP-glo IC_50_ / ULK1 NanoBRET IC_50_** | **cogP (red ≥ 4)** |
| --- | --- | --- |
| SBI-0206965 | 6 | 2.9 |
| 19 | 8 | 3.0 |
| 7 | 9 | 3.8 |
| 10 | 16 | 2.4 |
| 21 | 20 | 3.4 |
| 3 | 23 | 3.3 |
| SBI-5147 (5) | 24 | 4.1 |
| SBI-0757455 | 25 | 4.2 |
| 15 | 30 | 2.5 |
| SBI-7501 (2) | 42 | 3.9 |
| 1 | 47 | 3.8 |
| 6 | 53 | 3.8 |
| 14 | 75 | 4.1 |
| 13 | 76 | 2.8 |
| 20 | 116 | 4.7 |
| 11 | 137 | 4.2 |

Potency ratios were calculated between the compound potency in ULK1 ADP-glo versus ULK1 NanoBRET (See Table 1). Shown are all compounds with ADP-glo potencies < 200 nM and NanoBRET potencies < 5 µM. Calculated LogP values were estimated using ChemDraw. Highlighted in red are cLogP values > 4.

**Table S2. ULK1 & ULK2 ADP-Glo and NanoBRET IC_50_ values (nM).**

| **Compound ID** | **ULK1 ADP-Glo** | **ULK2 ADP-Glo** | **ULK1 NanoBRET** | **ULK2 NanoBRET** |
| --- | --- | --- | --- | --- |
| SBI-0206965 | 130 ± 20 | 2448 ± 298 | 785 ± 26 | 4695 ± 110 |
| SBP-7455 | 13 ± 2 | 476 ± 21 | 328 ± 43 | 1319 ± 202 |
| 1 | 5 ± 1 | 64 ± 22 | 236 ± 60 | 172 ± 33 |
| SBP-7501 (2) | 4 ± 0 | 118 ± 34 | 167 ± 31 | 174 ± 13 |
| 3 | 8 ± 0 | 704 ± 63 | 186 ± 5 | 1103 ± 454 |
| SBP-5147 (5) | 2 ± 0 | 53 ± 41 | 47 ± 11 | 17± 5 |
| 6 | 13 ± 3 | 217 ± 108 | 685 ± 41 | 1467 ± 261 |
| 7 | 31 ± 4 | 896 ± 92 | 291 ± 28 | 1583 ± 325 |
| 14 | 6 ± 5 | 464 ± 90 | 448 ± 142 | 1820 ± 599 |
| 15 | 20 ± 3 | 980 ± 62 | 605 ± 100 | 875 ± 131 |
| 19 | 16 ± 4 | 480 ± 65 | 132 ± 10 | 77 ± 13 |
| 21 | 5 ± 3 | 50 ± 12 | 100 ± 3 | 44 ± 10 |

Biochemical binding affinity (ADP-Glo) and intracellular target engagement (NanoBRET) of selected ULK1/2 inhibitors against ULK1 and ULK2. IC_50_ values indicate inhibitory potency, and results represent the means ± SD of three independent experiments performed in triplicate (ULK1/2 ADP-Glo, ULK2 NanoBRET) or duplicate (ULK1 NanoBRET).

**Table S3. ULK2 Protein Thermal Shift (PTS) of selected compounds.**

| **Compound ID** | **ULK2 PTS (ºC)** |
| --- | --- |
| SBI-0206965 | 8.0 ± 0.3 |
| SBP-7455 | 9.5 ± 0.2 |
| SBP-7501 (2) | 9.5 ± 0.1 |
| 3 | 3.0 ± 1.3 |
| SBP-5147 (5) | 14.0 ± 0.1 |
| 6 | 7.0 ± 0.7 |
| 7 | 2.0 ± 0.8 |
| 14 | 4.0 ± 0.1 |
| 15 | 8.0 ± 0.1 |
| 19 | 7.5 ± 0.0 |
| 21 | 8.0 ± 0.6 |

Protein thermal shift data of selected compounds against ULK2 kinase domain. Data represent the means ± SD of three independent experiments performed in triplicate.

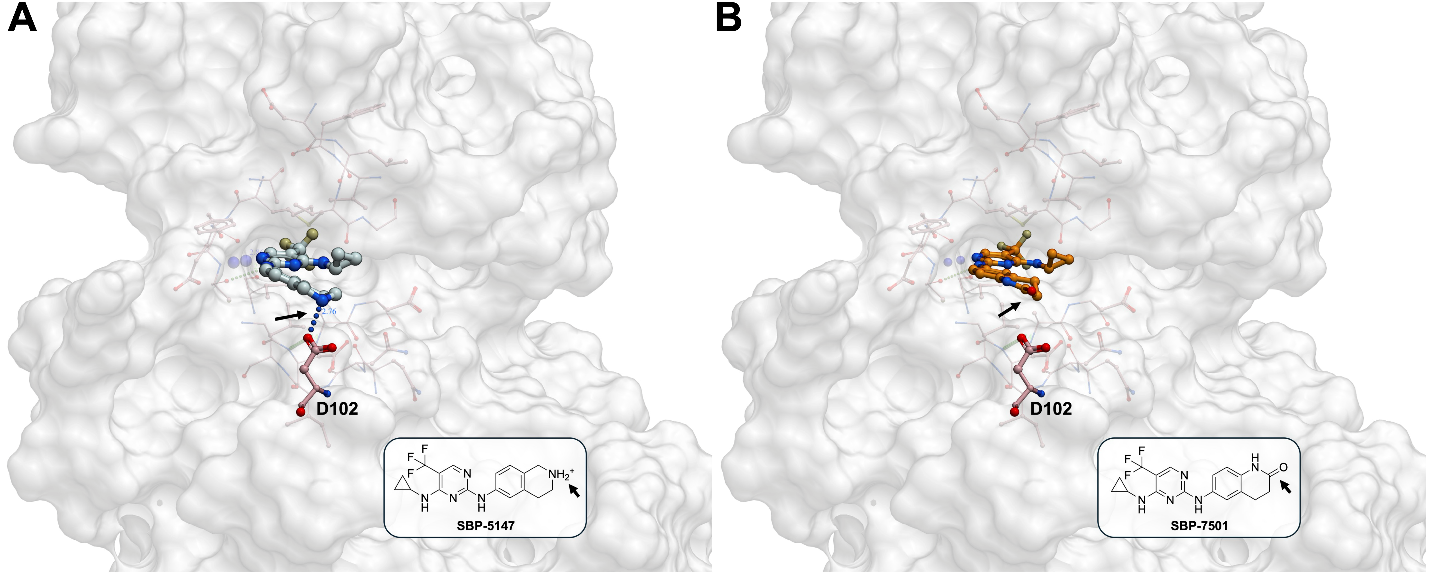

**Figure S3. Interaction of ULK1 inhibitors with the Asp102 side chain. (A)** Crystal structure of SBP-5147 bound to ULK1 (PDB ID: 9SE8). The ionic interaction between the protonated secondary amine of the 1,2,3,4-tetrahydroisoquinoline moiety and the Asp102 side chain is indicated by an arrow; dotted lines represent hydrogen bonds. **(B)** Flexible ligand docking of SBP-7501 into the ULK1 structure (PDB ID: 9SE8) using ICM Pro (Molsoft, LLC). The compound occupies a position similar to SBP-5147 in the crystal structure; however, the 3,4-dihydroquinolinone moiety (arrow) cannot directly engage the Asp side chain, potentially accounting for SBP-7501’s reduced potency. Similar results were obtained for SBP-7501 with ULK2.

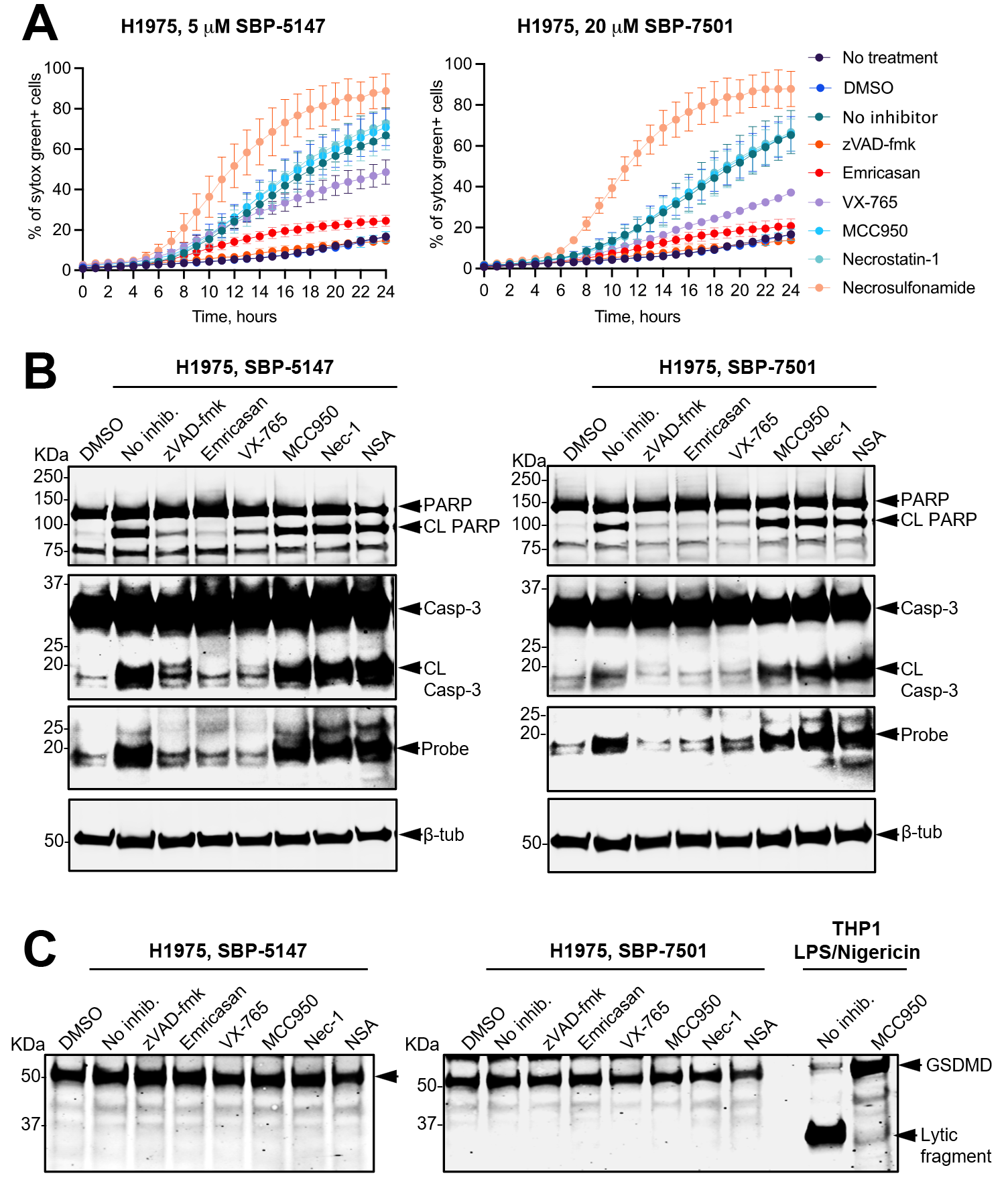

**Figure S4. ULK1/2 inhibitors induce apoptosis in H1975 cells.** (**A**) Cell death assay. H1975 cells were treated for 1 h with inhibitors that target caspases or other cell death signaling proteins (zVAD-fmk, Emricasan, MCC950, Necrosulfonamide, VX-765, and Necrostatin-1). Subsequently, cells were treated with **SBP-5147**, **SBP-7501**, or DMSO and imaged for 24 h. Total cell numbers were calculated by Hoechst staining and death cells were detected by sytox green staining. The graph shows the means ± SD of three independent experiments. (**B**) Apoptosis protein markers were detected by immunoblotting. Cells were treated as described above (panel A) and lysates were collected after 8 h. (**C**) Detection of the pyroptosis effector protein, gasdermin D. THP-1 cells were treated with LPS for 4 h followed by Nigericin for 2 h, with or without MCC950. Western blots are representative of two independent experiments. The probe (biotin-ahx-DEVD-AOMK), which reveals active caspase and cleaved proteins, is identified by “CL”.

**Table S4. Data collection and refinement statistics for ULK1 and ULK2 structures.**

| **Complex** | **ULK1 SBP-5147** | **ULK2 SBP-5147** |
| --- | --- | --- |
| **PDB accession code** | **9SE8** | **9SE9** |
| **Data Collection** |  |  |
| Resolution^a^ (Å) | 19.8-2.00 (2.07-2.00) | 63.7-1.97 (2.04-1.97) |
| Space group | *P*4_3_2_1_2 | *P*2_1_ |
| Cell dimensions | a = 74.2, b = 74.2, c = 220.7 Å | a = 76.6, b = 78.1, c = 94.2 Å |
|  | α, β, γ = 90.0° | α, γ = 90.0˚, β = 98.8˚ |
| No. unique reflections^a^ | 42,703 (4,097) | 77,687 (7,585) |
| Completeness^a^ (%) | 99.9 (100) | 100 (100) |
| I/σI^a^ | 11.4 (2.1) | 13.1 (2.0) |
| R_merge_^a^ | 0.107 (0.962) | 0.078 (0.919) |
| CC (1/2) | 0.998 (0.725) | 0.999 (0.669) |
| Multiplicity^a^ | 7.2 (7.3) | 6.4 (6.3) |
| **Refinement** |  |  |
| No. atoms in refinement (P/L/O)^b^ | 4,401 / 50 / 417 | 8,409 / 100 / 451 |
| Mean B value (Å^2^) | 37.3 | 48.9 |
| R_fact_ (%) | 18.3 | 19.3 |
| R_free_ (%) | 22.8 | 23.0 |
| rms deviation bond^c^ (Å) | 0.013 | 0.013 |
| rms deviation angle^c^ (°) | 1.38 | 1.35 |

^a^ Values in parentheses are the statistics for the highest-resolution shells.

^b^ P/L/O indicates protein, ligand molecules present in the active sites, and other (water, solvent, ion) molecules, respectively.

^c^ rms indicates root-mean-square.
